# Endogenous retroelements promote tolerance to dietary antigens

**DOI:** 10.1101/2025.09.09.675192

**Authors:** Claudia A. Rivera, Eduard Ansaldo, Verena M Link, Siddharth R Krishnamurthy, Ana Teijeiro, Cihan Oguz, Daniel Yong, Yasmine Belkaid

## Abstract

Retroelements are transposable elements that represent a significant portion of eukaryotic genomes. Here, we show that constitutive expression of endogenous retroelements play a key regulatory role in the acquisition of food tolerance. Specifically, inhibition of retroelement reverse transcription abolishes tolerance to dietary antigens and promotes allergic responses. This phenomenon is associated with impaired regulatory T cell differentiation/accumulation and altered dendritic cell tolerogenic function. Mechanistically, innate sensing of retroelement-derived cDNA via cGAS/STING within gut epithelial cells promotes a local tolerogenic milieu. Thus, within the gut, immune reactivity to retroelements act as a local tonic signal required for regulatory T cell induction and differentiation, thereby preventing allergic responses to food. Collectively, these findings uncover retroelements as key regulatory elements and essential allies in maintaining immune tolerance.

## Main Text

As multicellular organisms evolved in microbe- and nutrient-diverse environments, the gut became a critical interface where the immune system had to balance defense and tolerance. The ability to suppress immune responses to harmless antigens conferred a selective advantage by preserving tissue integrity and ensuring efficient nutrient absorption. Failure of these regulatory mechanisms is associated with aberrant reactivity to oral antigens, a phenomenon underlying allergic responses. Over the past few decades, the global incidence of food allergies has risen at an accelerated pace (*1*, *2*), highlighting the urgent need to identify the critical pathways underlying both adaptive and maladaptive immune responses to oral antigens.

Within the gut, the evolutionary pressure to maintain tolerance to continual antigenic exposure shaped the development of specialized immune structures and regulatory pathways. Previous work by us and others has shown that within the gastrointestinal tract, epithelial cells and dendritic cells play crucial roles in shaping immunoregulatory responses. More specifically, local dendritic cells (DCs), through the production of various factors including TGF-β and retinoic acid (RA), promote the differentiation of food antigen-specific regulatory T cells (T_regs_), which in turn suppress inappropriate immune responses to dietary or commensal derived antigens (*3–5*). Upstream of these responses, the nature of the signals involved in the maintenance of a tolerogenic milieu remains an active area of investigation.

Endogenous retroelements (EREs) make up a substantial portion of the mammalian genome— about 42% in humans and 37% in mice (*6*, *7*). These retrotransposable elements include endogenous retroviruses (ERVs)—remnants of ancient viral infections—and long interspersed nuclear elements-1 (LINE-1) (*8*, *9*). Both ERVs and LINEs elements can retain functional open reading frames and are actively transcribed into RNA, some of which are reverse transcribed into cDNA by encoded reverse transcriptase (RT) enzymes (*10*). These retroelement-derived products can activate innate immune sensors that detect cytosolic RNA and DNA, triggering downstream antiviral responses, including the production of type I interferons (IFN-I) (*11*).

Emerging evidence supports the concept that, through their ability to induce both innate and adaptive immune responses, these ancient and intrinsic components of our genome play a fundamental role in regulating host physiology and immunity. Indeed, endogenous retroelements have been implicated in various pathological processes, including cancer, infection, and autoimmunity (*12–15*). Under steady-state conditions, we previously demonstrated that ERE expression by keratinocytes in the skin sets the tissue activation threshold in an IFN-I–dependent manner (*16*). Whether such a phenomenon occurs at other barrier sites, and to which extent EREs contribute to gut homeostatic function has not yet been addressed.

Of note, type I interferons have been shown to promote T_reg_ induction and previous work supported a role for IFN-I in the development of oral tolerance (*17*, *18*). In this context, it is intriguing to speculate that one potential source of IFN-I tonic signals may be controlled by ERE. More generally, the vast number of endogenous retroelements, along with their retained expression, represents a significant yet poorly explored potential for interaction with host physiology and for shaping local physiological functions. Here, we explore the hypothesis that within the gut, EREs may act as keystone regulators in tissue-specific settings, co-opting host mechanisms to establish vital immunoregulatory responses.

## Results

### Endogenous retroelement reverse transcription promotes tolerance to dietary antigens independently of the microbiota

To assess whether retroelement expression could contribute to oral tolerance, we employed a model where reactivity to food antigen is assessed via intradermal challenge and associated delayed-type hypersensitivity reaction (DTH) (**Fig. 1A**). Using this model, we inhibited cDNA-sensing of retroelements during the tolerization phase by treating mice with a combination of reverse transcriptase inhibitors (anti-RT), tenofovir, and emtricitabine, which obstruct nascent cDNA synthesis (*16*). Consistent with previous reports (*19*, *20*), mice tolerized with the model antigen Ovalbumin (OVA) in drinking water developed mild to no-response following intradermal OVA challenge (**Fig. 1B, C, D**). In contrast, mice non-tolerized developed severe skin inflammation associated with recruitment of neutrophils and basophils (**Fig. 1B, C, D**). Inhibition of retroelement cDNA synthesis was associated with a breakdown of oral tolerance as evidenced by skin swelling, and increased neutrophils and basophil recruitment after challenge in a manner comparable to non-tolerized mice (**Fig. 1B, C, D**). OVA-specific IgG1 serum antibodies followed a similar pattern with increased antigen-specific IgG1 levels in tolerized anti-RT treated mice, showing similar levels as non-tolerized mice (**Fig. 1E and fig. S1A**). Taken together, these results propose that retroelement reverse transcription controls, at least partially, the induction of tolerance to dietary antigens.

**Figure 1.**
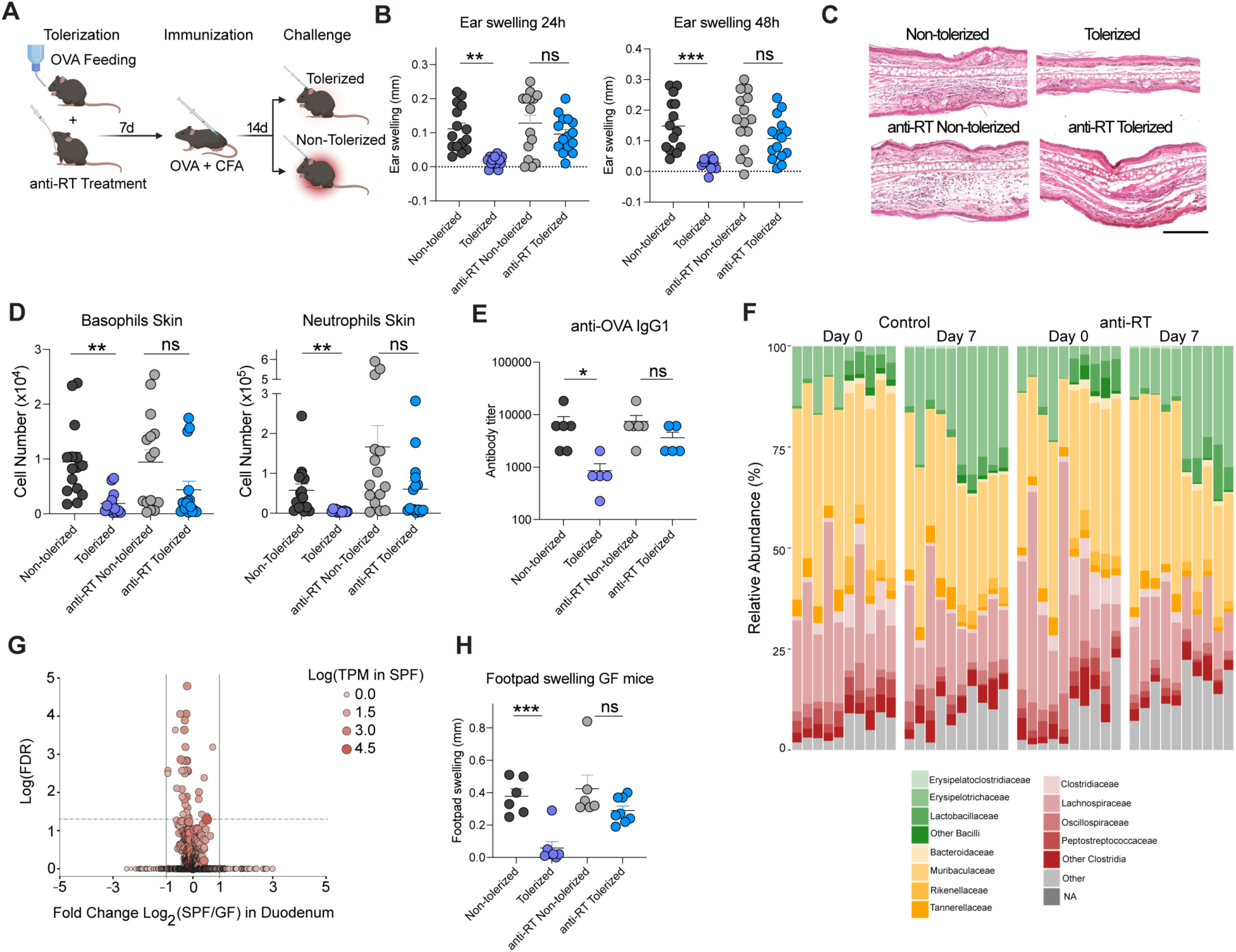
Reverse transcription of endogenous retroelements promotes tolerance to food antigens independent of the microbiota. (**A**) Scheme of DTH-induced responses in mice after OVA administration in drinking water and antiretroviral or control treatment during tolerization. (**B**) Mice were intradermally challenged with OVA in the ear, and ear swelling was measured after 24 and 48 hours. (**C**) Representative images of H&E staining of transverse sections from ear pinnae after DTH and intradermal OVA challenge. Scale bar 5 μM. (**D**) Absolute numbers of basophils and neutrophils infiltration after 48h of intradermal challenge analyzed by flow cytometry. (**E**) OVA-specific IgG1 antibodies analyzed by ELISA in sera samples obtained 48h post ear challenge in the DTH model. Graph depicts 1 independent experiment representative of 3 independent experiments. (**F**) Relative abundance of taxonomic family in the gut microbiota of mice before and after anti-RT or control treatment, analyzed by 16S rRNA sequencing. (**G**) Volcano plot of bulk RNA-seq analysis of retroelement loci in gut epithelial cells from SPF or Germ-Free (GF) mice. (**H**) DTH responses were induced in GF mice following a protocol similar to (A). Challenge was performed in the hind footpad and footpad swelling was analyzed 48h after challenge. Data are representative of three independent experiments. Each dot represents an individual mouse. Numbers in flow plots indicate mean ± SEM. For (B) and (H), one-way ANOVA with Tukey’s multiple comparisons test was used; for (D) and (E), Kruskal-Wallis with Dunn’s multiple comparisons test was used. * p < 0.05; ** p < 0.01; *** p < 0.001; **** p < 0.0001; ns, not significant.

We previously showed that retroelement sensing was required to orchestrate microbiota immune interactions within the skin (*16*). To address whether a microbiota-ERE dialogue could contribute to oral tolerance induction, we first analyzed the microbiota composition following inhibition of retroelement cDNA synthesis. Anti-RT treatment for 7 days was not associated with significant alteration in gut microbiota composition, abundance and diversity (**Fig. 1F and fig. S1B**). Further, analysis of retroelement expression in duodenal epithelial cells from germ-free (GF) versus specific-pathogen free (SPF) untreated mice, showed little to no changes (**Fig. 1G**). Finally, we tested the effect of retroelements in the induction of oral tolerance in mice devoid of microbiota. Inhibition of RT during tolerization in GF mice was associated with a breakdown of oral tolerance in a manner comparable to what was seen in the presence of the microbiota, supporting the idea that sensing of the microbiota or microbiota alteration is not associated with retroelement control of food tolerance (**Fig. 1H**).

### Endogenous retroelement reverse transcription promotes small intestine regulatory T cell differentiation and effector functions

To assess the impact of retroelement reverse transcription inhibition on gut immunity, we performed immune profiling of various immune populations within the small intestine, large intestine, and mesenteric lymph nodes (mLNs) after 7 days of anti-RT treatment. Myeloid and lymphoid populations were not impacted within the colon and mLN (**fig. S2A, S2B**). On the other hand, in the small intestine lymphoid compartment, T_regs_ were specifically decreased following anti-RT treatment, including those expressing GATA3 and RORγT (**Fig. 2A, B**), which have been previously associated with tolerogenic responses to food and the microbiota (*21–23*).

**Figure 2.**
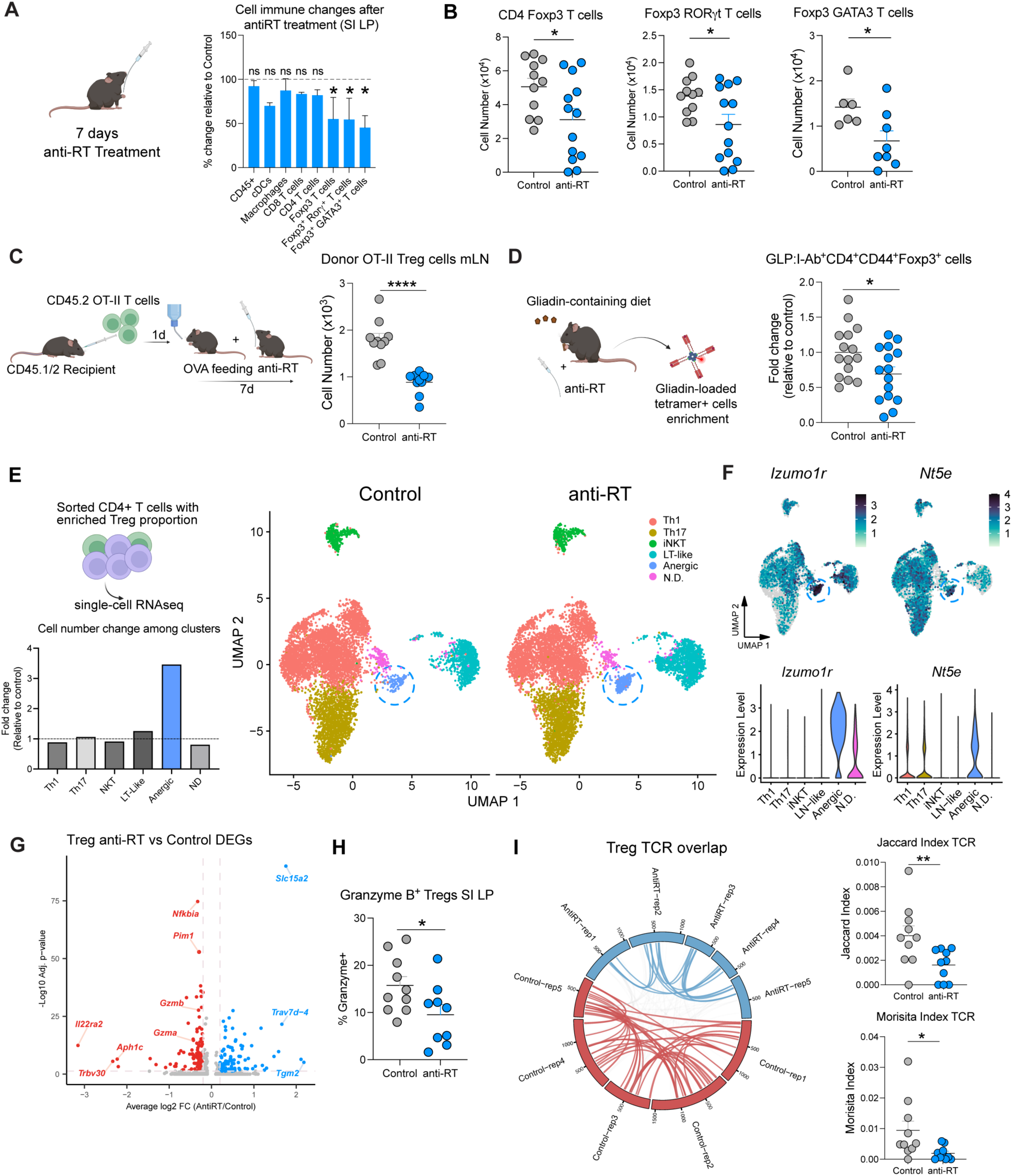
Endogenous retroelement reverse transcription supports gut T_reg_ effector functions. (**A**) Percentage change relative to control in absolute numbers of the indicated immune populations in small intestine lamina propria after 7 days of antiretroviral treatment. (**B**) Live CD45^+^ CD90.2^+^ TCRβ^+^ CD4^+^ Foxp3^+^ cells, live CD45^+^ CD90.2^+^ TCRβ^+^ CD4^+^ Foxp3^+^ RORγT^+^ cells and live CD45^+^ CD90.2^+^ TCRβ^+^ CD4^+^ Foxp3^+^ GATA3^+^ cells absolute number quantification by flow cytometry analysis. (**C**) (Left) Congenic CD45.1/2 recipient mice were adoptively transferred with CD45.2 OT-II OVA specific transgenic naïve CD4^+^ T cells purified from the spleen. Mice were provided OVA in drinking water and orally gavaged with anti-reverse transcriptase (anti-RT) treatment or control vehicle for 7 days. (Right) Absolute number of Live CD45.2^+^ CD90.2^+^ TCRβ^+^ CD4^+^ Foxp3^+^ cells analyzed by flow cytometry in mesenteric lymph nodes (mLN) after OT-II adoptive transfer. (**D**) (Left) Mice were fed a Gliadin-containing diet for 7 days and orally gavaged with anti-RT treatment or control vehicle for 7 days. Gliadin-loaded tetramer^+^ cell numbers were analyzed by flow cytometry in pooled secondary lymphoid organs (SLOs: mLNs, Peyer’s patches, spleen and hepatic LNs). Graph depicts fold change of Glp:I-A^b+^ tetramer^+^ CD4^+^ CD44^+^ Foxp3^+^ cells relative to control mice. (**E**) CD4^+^ T cells with an enriched proportion of Foxp3^+^ T cells sorted from SI LP from Foxp3 GFP^+^ reporter mice, treated with anti-RT or control, were analyzed by scRNA-seq. Fold change of cell abundance per different scRNA-seq clusters (excluding Foxp3^high^ T cells) after 7 days of antiretroviral treatment relative to control (Lower panel). UMAP projection plot shows major CD4^+^ T cell clusters. LT-like, lymphoid-tissue like; N.D., not determined. Dotted lines highlighting the anergic cluster (Right panel). (**F**) Expression of the anergic markers *Izumo1r* (FR4) and *Nt5e* (CD73) among different clusters in the UMAP representation of CD4^+^ T cell clusters (Top) or violin plots (Bottom) among different CD4^+^ T cell clusters. (**G**) Volcano plot of differentially expressed genes in T_regs_ after antiretroviral treatment. Red denotes downregulated and blue upregulated genes by anti-RT treatment. (**H**) Percentage of Granzyme B^+^ T_regs_ analyzed by flow cytometry after antiretroviral treatment. (**I**) Circos plot of T_reg_ T cell Receptor (TCR) analysis comparing control vs anti-reverse transcriptase treated mice. Each segment represents a mouse. Links between segments represent shared TCR between mice and colored links represent shared TCR between mice under the same treatment (Left). Jaccard and Morisita indexes of T_reg_ TCRs where each dot represents TCR overlap between mice under the same treatment (Right). For flow cytometry analyses, data are representative of at least two independent experiments. Each dot represents an individual mouse. Numbers in flow plots indicate mean ± SEM. For (A), (B), (C), (D), (H) two-tailed unpaired Student’s t-test was used. For (I), Mann-Whitney test was used. * p < 0.05; ** p < 0.01; **** p < 0.0001; ns, not significant.

We and others previously showed that T_reg_ induction played a fundamental role in oral tolerance to food antigens (*4*, *5*, *20*, *24*). To test whether retroelement sensing could impact food antigen-specific T_reg_ induction, OVA specific transgenic OT-II CD4^+^ T cells were transferred to naïve syngeneic recipients prior to oral administration of OVA in drinking water (*24*), in the context of antiretroviral treatment (**Fig. 2C**). In this model, T_reg_ induction is first detected within the mesenteric lymph nodes (mLN), prior to accumulation in the small intestine lamina propria (*19*). Anti-RT treatment at the time of OVA exposure significantly reduced the differentiation and/or accumulation of T_regs_ within the mLNs (**Fig. 2C**). We next employed another model of T_reg_ induction associated with oral responses to the wheat protein gliadin (*25*). In this model, gliadin-specific cells can be detected via recently reported Glp:I-A^b^ tetramers (*25*). Mice fed with a gliadin-containing diet showed reduced accumulation of tetramer-binding T_regs_ within secondary lymphoid organs following anti-RT treatment compared to control mice (**Fig. 2D, fig. S2C**). Thus, retroelement reverse transcription controls food antigen-specific responses and food-specific T_reg_ induction.

To assess the potential impact of antiretroviral treatment on T_reg_ phenotype and function, we next performed a droplet-based 5’ single-cell RNA sequencing (scRNA-seq) of sorted CD4^+^ T cells with an enriched proportion of Foxp3^+^ T cells (1:1 ratio) to allow capture of T_reg_ precursors. Analysis of conventional CD4^+^ T cells (Foxp3^-^) showed that cluster proportions of Th1 or Th17 cells were unaffected (**Fig. 2E**). On the other hand, a cluster of CD4^+^ T cells with high expression of Izumo1r (FR4) and Nt5e (CD73), two classical anergic markers, was expanded post-treatment (**Fig. 2E, F, fig. S2D**). Of interest, recent work identified this population as precursors of food antigen specific T_regs_ (FR4^+^Th^lin-^ cells) (*25*). Because of the decrease in absolute numbers of mature T_reg_ post treatment, such an increase in T_reg_ precursors supports the idea that anti RT treatment may be associated with arrested/delayed T_reg_ development.

Within mature T_reg_ cells (**fig. S2E**), several canonical effector T_reg_ genes were downregulated following antiretroviral treatment (**Fig. 2G**). Specifically, *Gzma* and *Gzmb*, showed lower expression compared to control condition, which we further validated by flow cytometry (**Fig. 2H**). Granzyme B has been previously reported to contribute to T_reg_ effector functions in the context of oral tolerance (*26*). Interestingly, *Nfkbia*, encoding IkBa, directly interacts with RelA, a NF-κB subunit that regulates T_reg_ effector differentiation and survival (*27*). Consistent with our phenotype, T_reg_ downregulated the expression of *Nfkbia* after antiretroviral treatment. Inhibition of reverse transcription also downregulated expression of *Pim1*, a TNF transducer, reported as a marker of non-lymphoid tissue T_regs_ (*28*) (**Fig. 2G**). The TNF receptor superfamily-NF-κB axis has also been shown to promote the maintenance of effector T_regs_ in non-lymphoid tissues (*29*). Thus, our results support the idea of a reduced fitness and/or maturation of T_reg_ cells post anti-RT treatment.

Previous work revealed that T_regs_ undergo clonal TCR selection after encountering dietary protein antigens (*26*). Such clonality was significantly reduced post antiretroviral treatment as evidenced by decrease of TCR overlap among mice (**Fig. 2I**). Of note, inhibition of retroelement cDNA synthesis did not affect TCR clonality in CD4^+^ conventional T cells (**fig. S2F**). Thus, our results propose that retroelement reverse transcription promotes T_reg_ effector differentiation, maturation and clonal expansion.

### Retroelement reverse transcription promotes dendritic cell homeostasis and tolerogenic function

Within the gastrointestinal tract, various antigen presenting cells (APCs) contribute to T_reg_ induction (*3–5*). In adult mice, conventional dendritic cells (cDCs) are one of the main APCs presenting dietary antigens. Two subsets have been shown to contribute to tolerogenic responses: CD103^+^ conventional DCs type 1 (cDC1s) induce food antigen-specific T_regs_, and CD103^+/-^ conventional DCs type 2 (cDC2s) likely contribute to oral tolerance via induction of T_reg_ precursors (FR4^+^Th^lin-^) (*3*). This process is also controlled by the ability of gut cDCs to produce large amounts of both retinoic acid (RA) and TGF-β (*4*, *5*, *30*, *31*). Indeed, following anti-retroviral treatment, both RA activity and TGF-β production were significantly impacted in CD103^+^ cDC2s and cDC1s from the small intestine (**Fig. 3A, B**).

**Figure 3.**
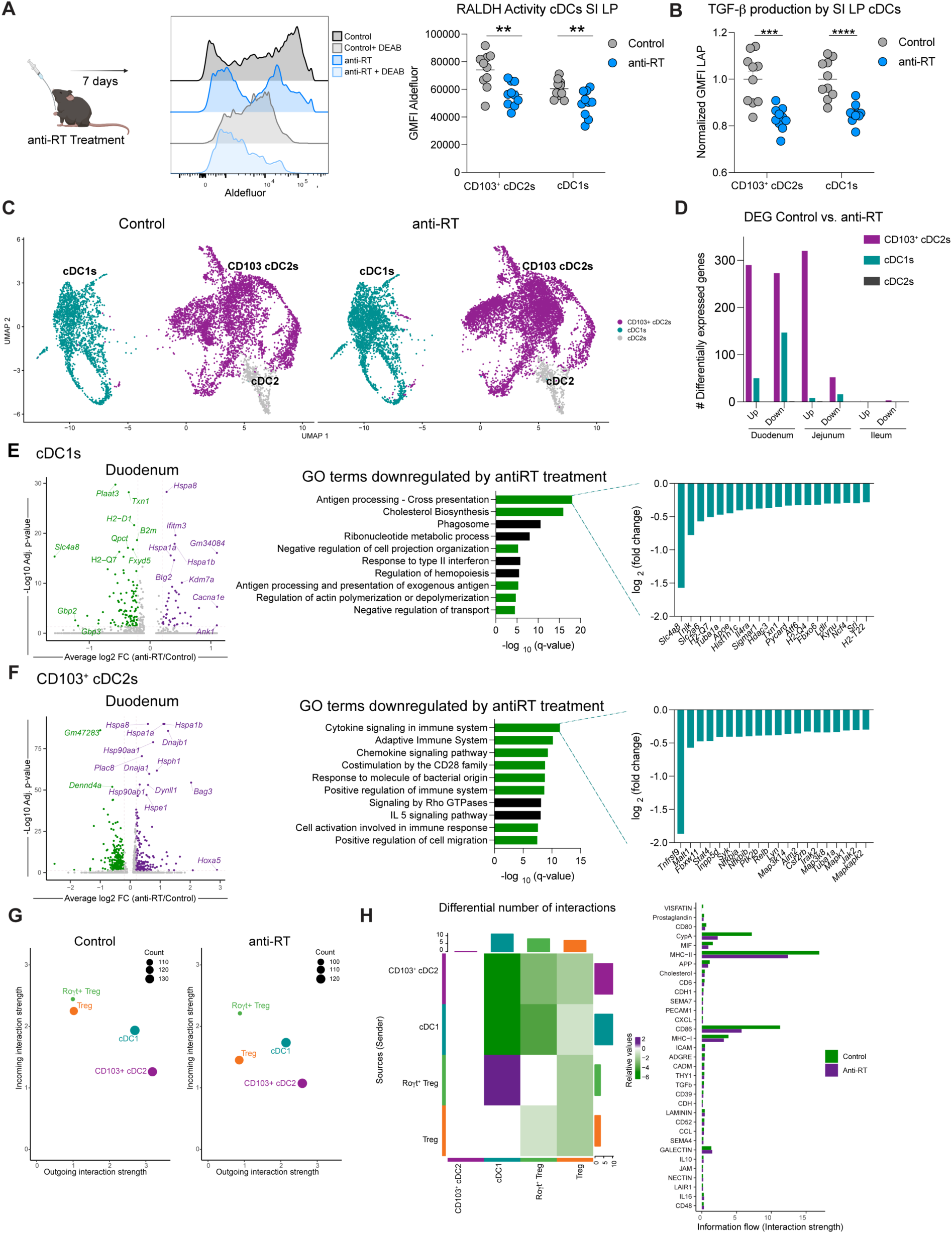
Retroelement reverse transcription impacts intestinal dendritic cell homeostasis. (**A**) Retinoic acid/aldehyde dehydrogenase activity was analyzed by flow cytometry after anti-RT treatment using Aldefluor kit. Plot depicts geometric mean fluorescence intensity (GMFI) of aldefluor in small intestine CD103^+^ cDC2s and cDC1s. (**B**) Normalized GMFI of LAP (TGF-β1) analyzed by flow cytometry in small intestine CD103^+^ cDC2s and cDC1s after cytokine stimulation. (**C**) Live CD45^+^ Lineage^-^ (TCRβ^-^ TCRγδ^-^ NK1.1^-^ CD45R^-^) CD64^-^ MHC-II^high^ CD11c^high^ cells from small intestine were analyzed by scRNA-seq. UMAP projection of dendritic cells between control or antiretroviral treated mice. (**D**) Number of differentially expressed genes (DEG) among different dendritic cell populations in duodenum, jejunum and ileum after anti-RT treatment. (**E**) Volcano plot of DEGs in duodenal cDC1s after inhibition of reverse transcriptase treatment. Purple denotes upregulated and green downregulated by anti-RT treatment. (Left). GO terms in cDC1s downregulated after anti-RT treatment (Middle). Downregulated genes related to antigen processing-cross presentation pathway are shown (Right). (**F**) Volcano plot of DEGs in duodenal CD103^+^ cDC2s after inhibition of reverse transcriptase treatment. Purple denotes upregulated and green downregulated by anti-RT treatment. (Left). GO terms in CD103^+^ cDC2s downregulated after anti-RT treatment (Middle). Downregulated genes related to cytokine signaling in immune system pathway are shown (Right). (**G**) Scatter plot showing the total outgoing (ligand) and incoming (receptor) signals (interaction strength) for T_reg_, RORγt^+^ T_reg_, and duodenal cDC1s and CD103^+^ cDC2s, identified using CellChat. Dot size is proportional to the number of interactions. (**H**) Heatmap depicting the differential number of anti-RT treatment ligand-receptor interactions relative to control state across all cell types. Bar height represents the degree of change in terms of the number of interactions between the two conditions: top bar plot represents sum of each column (incoming signaling: receiver), right bar plot represents sum of each row (outgoing signaling: sender) (left). Comparison of specific pathways between control and anti-RT treatment based on the differential interaction strengths of pathway-specific ligand-receptor interactions (right). For flow cytometry analyses, data are representative of at least two independent experiments. Each dot represents an individual mouse. For (A) and (B) multiple two-tailed unpaired Student’s t-test was used. Numbers in flow plots indicate mean ± SEM. ** p < 0.01; *** p < 0.001; **** p < 0.0001.

We next performed a scRNA-seq on sorted dendritic cells (identified as MHC-II^high^ CD11c^high^) isolated from the duodenum, jejunum, and ileum (**Fig. 3C and fig. S3A**). Strikingly, most changes in gene expression were found in DCs from the upper part of the small intestine (duodenum) where food is first seen within the small intestine. Fewer changes were observed in the jejunum and little or no variation observed in the ileum. Within the duodenum, anti-RT treatment modulated gene expression of both cDC1s and CD103^+^ cDC2, while CD103^-^ cDC2s were unaffected (**Fig. 3D, E, F** and **fig. S3B, S3C, S3D**). In the duodenum, cDC1s exhibited decreased expression of genes related to antigen processing and presentation, and downregulated pathways related to cell protrusions and migration (**Fig. 3E**). Within the CD103^+^ cDC2 population, inhibition of retroelement reverse transcription was associated with downregulation of genes related to cytokine and chemokine signaling, cell immune activation and cell migration (**Fig. 3F**). Ligand-receptor interaction analysis revealed that this antiretroviral treatment also affected cDC-T_reg_ interactions, by reducing the number and strength of signals sent from duodenal cDCs to regulatory T cells, both in terms of antigen presentation and co-stimulation (**Fig, 3G, H**). Thus, inhibition of reverse transcriptase has a significant impact on the activation and function of conventional DCs associated with the induction of tolerance to food antigen.

### Tonic endogenous retroelement expression is cell and tissue specific, and independent of exogenous factors

To determine where retroelement expression was needed to set the tone required for oral tolerance, we next assessed retroelement expression by gut epithelial cells, cDC1s, and CD103^+^ cDC2s at steady state and under various conditions. We first analyzed steady state ERE expression among cell subsets in the duodenum, the site where we observed the most significant changes in cDC function upon RT inhibition. Notably, cDC1s expressed a distinct and unique pattern of retroelement expression when compared to epithelial cells or CD103^+^ cDC2s, while the latter two populations showed similarities (**Fig. 4A, B**). These unique retroelement signatures included several LINE-1 associated elements, as well as LTRs (mostly ERVs), and Short Interspersed Nuclear Elements (SINE) (**Fig. 4A**). While CD103^+^ cDC2s showed higher variability among replicates, ERE expression patterns in cDC1s and epithelial cells remained more consistent. As we previously reported in the skin (*16*), anti-RT treatment did not impact ERE expression in both cDCs and epithelial cells (**fig. S4A**). At the protein level, ORF1p of Long-Interspersed Nuclear Element-1 (LINE-1) expression was higher in both cDC1s and epithelial cells from duodenum when compared to CD103^+^ cDC2s (**Fig. 4C**). Cell surface expression of the envelope gene derived from endogenized Murine Leukemia Virus (MLV) was higher in epithelial cells compared to cDCs (**Fig. 4C**).

**Figure 4.**
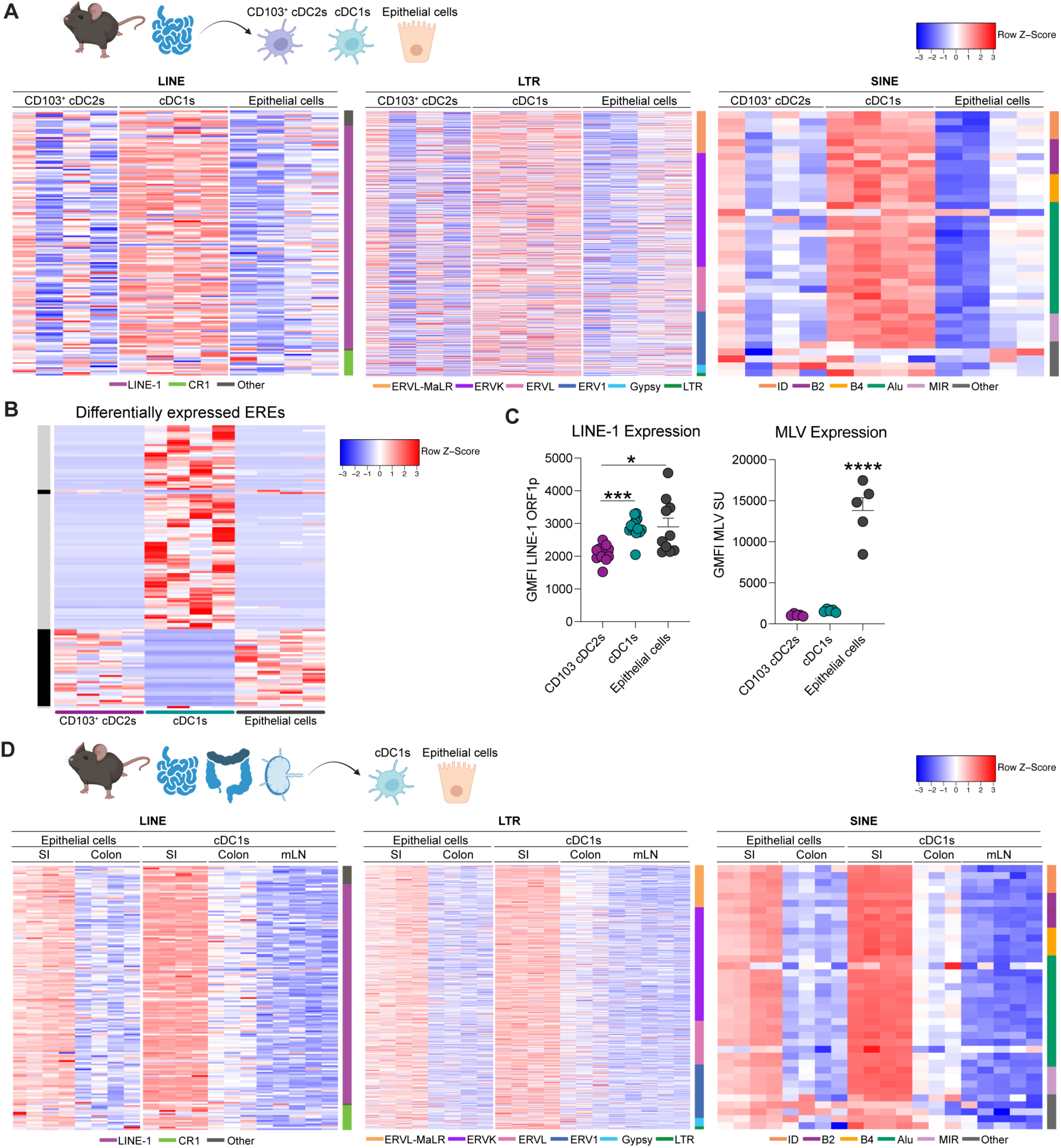
Tonic endogenous retroelement expression is tissue and cell specific. (**A**) Heatmap displaying row Z-score of ERE expression at the locus level from bulk RNA-seq in sorted epithelial cells, CD103^+^ cDC2s and cDC1s from duodenum. Long-Interspersed Nuclear Elements (LINE) (left), Long Terminal Repeats (LTR) (middle) and Short-Interspersed Nuclear Elements (SINE) (right). Families are listed on the bottom. CR1, chicken repeat 1; ERV, endogenous retrovirus; MaLR, mammalian apparent LTR retrotransposon; ID, identifier; MIR, mammalian-wide interspersed repeat. Multiple biological replicates are shown per cell type. (**B**) Heatmap depicting row Z-score of retroelement expression from bulk RNA-seq analysis from differentially expressed ERE at the chromosome level in same populations as in (A). Black depicts cDC1 vs. epithelial cells, and gray cDC1 vs. cDC2 comparison. (**C**) LINE-1 ORFp1 and MLV envelope protein displayed as GMFI in the same cell types as in (A) in duodenum analyzed at the protein level by flow cytometry. Left panel is representative of three independent experiments, and right panel depicts one independent experiment representative of at least 2 independent experiments. Each dot represents an individual mouse. One-way ANOVA with Tukey’s multiple comparisons test was used. * p < 0.05; *** p < 0.001. (**D**) Retroelement expression analyzed by bulk RNA-seq from sorted cDC1s from duodenum, colon and mesenteric lymph nodes (mLNs) and sorted epithelial cells from colon and duodenum. Heatmap displays row Z-score or differentially expressed ERE loci for the same families as in (B).

Previous work revealed that activation of cells via Toll-like receptors (TLRs) upregulated retroelement expression (*32*). Building on this, we aimed to determine whether endogenous retroelement expression is modulated by TLR signaling within the gut. To this end, we utilized TLR-deficient mice (Tlr2^−/−^ × Tlr4^−/−^ × Unc93b1) which lack all functional TLR signaling (*33*) and compared them to co-housed wild-type (WT) animals. Surprisingly, the absence of TLR function minimally impacted ERE expression across all cell types analyzed (**fig. S4B**).

Next, we examined whether dietary components could influence ERE expression in the gut. Mice were fed either a regular chow diet, an amino acid (AA) diet, or a casein-based diet. While the casein and AA diets are nutritionally equivalent, the AA diet contains only free amino acids, lacking intact protein antigens found in casein. On the other hand, the chow diet is a complex mix with a diverse composition of proteins and metabolites. Remarkably, retroelement expression remained consistent across all diets and cell types, regardless of nutritional content (**fig. S4C**). Furthermore, we compared retroelement expression in cDC1s and epithelial cells across various tissues (colon, small intestine, and mesenteric lymph nodes). Strikingly, ERE expression was higher in the small intestine for both cell types and across different ERE families (**Fig. 4D**). These findings highlight a remarkable partitioning of retroelements within the gut and a surprising stability of expression across diverse nutritional and innate stimulation settings. These results indicate that within the gut, ERE expression is constitutive, further supporting the concept that these genomic elements may be involved in canonical homeostatic functions.

### The cGAS-STING signaling pathway is required for the establishment of tolerance to dietary antigens

Reverse transcribed retroelement-derived cDNA can be sensed by different pattern recognition receptors (PRRs): TLR9 and the cGAS/STING pathway, the two main extranuclear DNA sensors (*6*, *8*, *34*, *35*). We next assessed the potential contribution of these pathways in the control of oral tolerance. To avoid compensatory responses (a phenomenon often observed in the context of homeostatic responses), we favored approaches associated with temporal pathway neutralization. First, we targeted TLR9 signaling by using inhibitory oligonucleotides (ODN), specifically during the tolerization phase of the DTH model. ODN2088 had no impact on the acquisition of oral tolerance, as evidenced by unaltered skin responses to OVA after tolerization (**fig. S5A**).

We previously reported that ERV cDNA was sensed by the cGAS-STING pathway in keratinocytes within the skin (*16*). To determine whether the cGAS-STING pathway could contribute to the induction of tolerance to dietary antigens, we utilized the pharmacological STING inhibitor H151 during DTH tolerization. H151 administration concomitantly with OVA feeding abolished oral tolerance in a manner comparable to anti-RT treatment at early time points (**Fig. 5A**). Of note, previous work demonstrated that gut explants released cGAMP, supporting the idea that this pathway is constitutively active within this compartment (*36*). Treatment of gut explants with anti RT for 24h, significantly reduced cGAMP production, supporting the role of the cGAS-STING pathway in the sensing of ERE gut activity (**Fig. 5B**).

**Figure 5.**
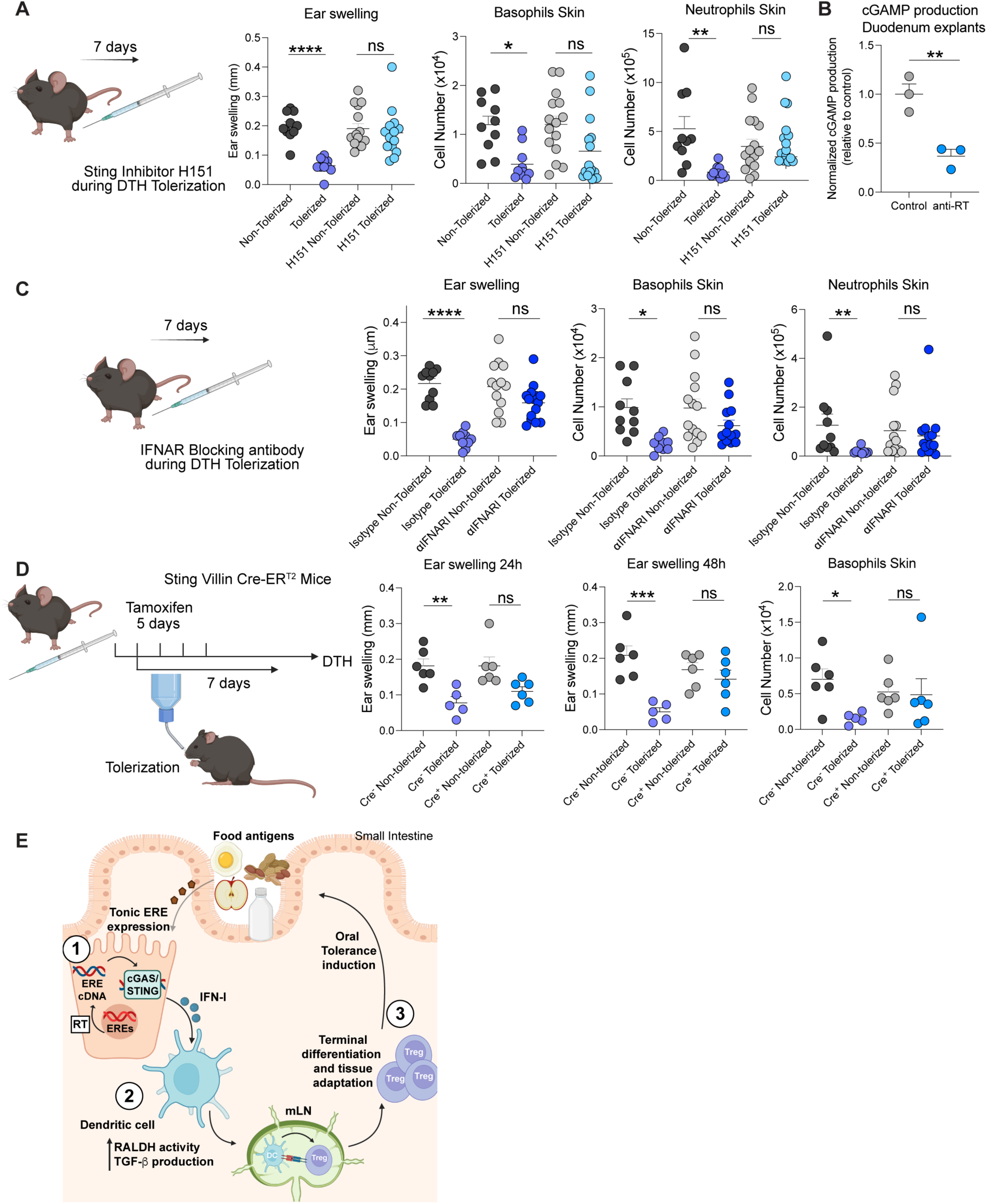
cGAS-STING signaling pathway and IFN-I are required for the establishment of oral tolerance. (**A**) STING was temporally blocked during DTH tolerization employing a small-molecule inhibitor (H151). Vehicle was injected as control. Ear swelling and infiltration of neutrophils and basophils were analyzed 24h later by flow cytometry in ear skin. (**B**) Gut explants were obtained from duodenum and cultured in vitro with antiretroviral treatment or control vehicle. cGAMP production was analyzed in supernatants after 24h by ELISA. (**C**) IFN-I responses were blocked during DTH tolerization by utilizing a blocking antibody to interferon alpha receptor 1 (IFNAR1). Mouse IgG1 isotype was administered as a control. Ear swelling and infiltration of neutrophils and basophils were analyzed after 24h by flow cytometry in skin. (**D**) STING expression was temporally deleted in gut epithelial cells by using tamoxifen right before and during DTH tolerization in Sting^flox/flox^ Villin Cre-ER^T2^ mouse models. Villin-Cre-^ERT2Cre-^ Sting^flox/flox^ or Villin-Cre-ER^T2Cre+^ Sting^flox/flox^ mice were used. Ear swelling was analyzed after 24h and 48h, and infiltration of basophils was analyzed after 48h by flow cytometry in skin. (**E**) Model of the role of EREs in controlling tolerance to dietary antigens. Data are representative of 3 independent experiments for B and C, and of 2 independent experiments, representative of 3 independent experiments for D. Each dot represents an individual mouse. Numbers in flow plots indicate mean ± SEM. For (A), (C) and (D), one-way ANOVA with Tukey’s multiple comparisons test or Kruskal-Wallis with Dunn’s multiple comparisons test were used. For (B) two-tailed unpaired Student’s t-test was used. * p < 0.05; ** p < 0.01; *** p < 0.001; **** p < 0.0001; ns, not significant.

The cGAS-STING pathway is associated with the release of either IFN-I or IFN-III (IFN-λ) (*37*–*39*). To assess whether IFN-I or -III responses contributed to the induction of food tolerance, mice were treated with a blocking antibody for the IFN-I receptor (interferon alpha receptor 1, IFNAR), a blocking antibody for IFN-λ 2/3, or isotype controls during OVA tolerization. Neutralization of IFN-I signaling but not IFN-III, recapitulated the phenotype observed following anti-RT treatment 24h post challenge (**Fig. 5C and fig. S5B**).

We next assessed in which specific compartment STING signaling was required. To this end, we temporarily deleted Sting in intestinal epithelial cells during tolerization using Sting^flox/flox^ mice in a Villin-Cre-ER^T2^ tamoxifen-inducible system. Tamoxifen was administered right before and during tolerization to control the time frame where the STING signaling was impaired. Notably, blocking STING signaling during tolerization impacted the induction of tolerance to food, recapitulating the phenotype observed after anti-RT treatment (**Fig. 5D**). Altogether, these results propose that cGAS/STING sensing of retroelement cDNA in gut epithelial cells controls the induction of tolerance to dietary antigens.

Collectively, our model proposes that innate sensing of retroelement products by gut epithelial cells modulates tolerogenic dendritic cell function and the differentiation of food-specific regulatory T cells. More specifically, our model supports the idea that constitutive endogenous retroelement expression by gut epithelial cells and reverse transcription of these elements, induce the cytosolic accumulation of ERE-derived cDNA. This cGAS/STING innate sensing of EREs by epithelial cells promotes type 1 IFN responses thereby impacting the status of activation of neighboring antigen presenting cells. Such phenomenon leads to an optimal tolerogenic state, supporting the induction and differentiation of regulatory T cells specific to dietary antigens. Following their migration back to the small intestine, oral antigen specific T_regs_ undergo further differentiation and tissue adaptation under the influence of ERE-activated epithelial cell /DC milieu (**Fig. 5E**).

## Discussion

Endogenous retroelements have coevolved with their hosts over millions of years, shaping and being shaped by host genome regulation and immune defenses. Emerging evidence from our group and others uncovered that immune reactivity to EREs can contribute to immune system function, enhanced response to infection and cancer immunosurveillance (*6*, *8*, *16*, *39*). Here, we uncover a canonical role for EREs as essential allies in maintaining immune tolerance in the gut.

The vast number of EREs in the genome represents an enormous yet poorly explored, potential for interaction with host physiology and disease processes. Previous work revealed that, in humans, ERE expression was highly cell-specific (*40*). Our findings expand on this by showing that, within the gut, ERE expression is not only cell-specific but also compartment-specific. Notably, basal ERE expression levels are significantly higher in the small intestine—particularly in the duodenum—compared to the colon, supporting the notion of environment-specific regulatory mechanisms. This differential expression may reflect the distinct functions of each compartment, including the small intestine’s role in maximal nutrient absorption. Furthermore, we found that within the small intestine, basal ERE expression is surprisingly independent of external stimuli such as diet, microbial ligands, or the microbiota. These findings support our model in which cell- and tissue-specific, constitutive expression of retroelements controls tissue threshold of activation and homeostatic function.

Here, we show that the cGAS/STING pathway in gut epithelial cells plays an essential role in promoting tolerogenic responses to dietary antigen. The cGAS/STING pathway plays numerous roles in gut immunity, ranging from induction of antiviral immunity upon microbiota activation, to contradictory roles in colorectal cancer (CRC) and bowel diseases (*41*, *42*). In the skin, we previously demonstrated that immunity to the microbiota was regulated by cGAS/STING-mediated sensing of retroelements in keratinocytes (*16*). In our present setting, the microbiota was not involved in regulating ERE expression by gut epithelial cells. This may be explained by the dramatic differences in microbial biomass between the gut and the skin and the need to limit overactivity to the microbiota within the gut environment. Whether a dominant role for ERE–cGAS/STING signaling is conserved across all epithelial cell types, including those less exposed to external stimuli, remains to be determined. Thus, it is plausible that innate sensing of retroelements is intricately linked to a wide range of homeostatic and pathological processes, particularly those driven by dysregulated epithelial responses.

Given the well-established regulatory role of type I IFNs in various biological processes, our observation provides a potential explanation for the origin of the tonic signals necessary for IFN-I production. Within the gut, the impact of IFN-I signaling is highly context-dependent, leading to both tolerogenic and pro-inflammatory responses. For example, IFN-I signaling regulates IL-10 production by tumor-associated regulatory T cells in colorectal cancer (*43*). In contrast, high levels of IFN-I induced upon viral infection can interfere with the promotion of tolerance to dietary antigens (*44*, *45*). Transient shutdown of regulatory processes is an important feature that permits the development of adaptive immunity when needed. Indeed, both our work and that of others showed that impaired T_reg_ induction and or function occurs in the context of infection and inflammatory processes (*45–47*). Whether ERE expression and in some cases aberrant expression of these elements contribute to these diverse outcomes remains to be determined.

Food allergy has risen dramatically in the last decades, up by ∼70% from 1999 to 2021 in children in the US (*48*, *49*). Reports from Europe and Australia show similar results, with rates of hospitalizations due to allergic reactions to food doubling in a decade (*1*). In this context, understanding key determinants and controllers of this process is essential to further develop new tools and therapies to counteract the food allergy epidemic. Moving forward, it would also be important to assess the potential impact of pre-exposure prophylaxis (PrEP) treatment or altered innate sensing of ERE on the etiology of allergic responses. Particularly, understanding the effect of PrEP in healthy individuals with Celiac disease predisposition or at risk of Th2-skewed allergic responses, could provide important insights into the role of retroelement reverse transcription in food allergy.

Our work uncovers retroelements as key regulatory elements and essential allies in maintaining immune tolerance in the gut. The observation that these dominant components of mammalian genome, once considered genomic parasites, have become allies in such a fundamental physiological function, reinforces the emerging concept that retroelements may be deeply intertwined with many aspects of host physiology. Collectively, this new understanding of the relationship between retroelements and host biology opens a promising field of investigation with enormous potential to unravel the logic of tissue physiology, and how disruption of these pathways may contribute to pathological states.

## Materials and methods

### Mice

Conventional Specific Pathogen Free (SPF) C57BL/6J CD45.1, C57BL/6J CD45.1/2, C57BL/6-Tg (Foxp3-HBEGF/EGFP)23.2Spar/Mmjax (Jax 032050) and C57BL/6-[Tg] TCR OT-II-[KO]Rag2 mice were obtained from the NIAID Taconic Exchange Program. Full TLR deficient (Tlr2^−/−^xTlr4^−/−^xUnc93b13d/3d) mice were obtained from the Gregory M Barton Laboratory (*33*) and cohoused at 3-5 weeks of age for 4 weeks prior to start experiments. B6.Cg-Tg(Vil1-cre/ERT2)23Syr/J (JAX 020282) and B6;SJL-*Sting1^tm1.1Camb^*/J (JAX 031670) were purchased from The Jackson Laboratory. Germ free C57BL/6NTac mice were bred and maintained in the NIAID Microbiome Program gnotobiotic animal facility. Age- and sex-matched 7-12 weeks of age mice were used in each experiment. Littermate controls were used for experiments involving mice bred and maintained at NIAID. Mice were provided a standard pelleted rodent diet (Chow diet) unless otherwise noted. For experiments with special diets, mice were fed Casein Diet (TD. 210528) or Amino acid diet (TD. 210529) since birth, purchased form Envigo Teklad Diets. All mice were bred and maintained at an American Association for the Accreditation of Laboratory Animal Care (AAALAC)–accredited animal facility and housed in accordance with procedures outlined in the Guide for the Care and Use of Laboratory Animals. All experiments were performed at NIAID under an animal study proposal (LHIM2E) approved by the NIAID Animal Care and Use Committee.

### Anti-reverse transcriptase treatment

Mice were administered a combination of Tenofovir disoproxil fumarate and Emtricitabine (ACROS Organics, 100mg/kg for Tenofovir and 60mg/kg for Emtricitabine) as previously reported (*16*). Both reverse transcriptase inhibitors were dissolved in water and administered daily by oral gavage in 200ul for 7 days. Water was administered as control for gavage feeding.

### Delayed-Type Hypersensitivity induction

Oral tolerance experiments were performed as described previously (*46*). Briefly, mice were tolerized with OVA (1.5%) in drinking water ad libitum for 6 days. Anti-RT was administered or not during tolerization by daily gavage for 7 days. One day after, mice were subcutaneously injected with a 1:1 emulsification with Complete Freund’s Adjuvant (CFA, Sigma-Aldrich). 14 days later, mice were challenged with 10μg OVA in PBS either by intradermal ear injection or in their hind footpad (for GF experiments, due to technical difficulties). Following intradermal challenge, ear or footpad thickness was measured with a digital caliper (Mitutoyo) 24 or 48 hours later. Ear and footpad swelling was reported as the difference between OVA-injected and un-injected ear/footpad thickness.

### OT-II T cells Adoptive transfer

CD4^+^ naïve T cells were purified from spleens and lymph nodes of OT-II-[KO]Rag2 mice using a magnetic enrichment kit (StemCell). 1x10^6^ cells in 100μl PBS were adoptively transfer by retro-orbital intravenous injection. 24h later, OVA was administered in the drinking water (1.5%) and anti-RT treatment was provided or not by oral gavage for 7 days. Mice were individually analyzed for the conversion of OT-II T cells into Foxp3^+^ CD4 T cells, employing CD45.1 and CD45.2 fluorophore-conjugated antibodies to discriminate donor/acceptor CD4^+^ T cells.

### In vivo treatments

TLR9 was blocked with a specific antagonist CpG ODN (ODN2088, Invivogen); mice were injected intraperitoneally (i.p.) with ODN2088 or ODN control, 10μg/mouse, every 3 days during the tolerization phase of DTH (total 3 injections). STING signaling blocking was performed by daily i.p. injection of the small molecule inhibitor H-151 (Invivogen); mice were injected with 200ul of 750nM H-151 in PBS 0.1% Tween-80 solution, or vehicle, daily for 7 days during DTH tolerization. IFN α/β receptor subunit 1 (IFNAR-I; clone MAR1-5A3, BioXCell) or IgG1 isotype control (clone MOPC-21, BioXCell) blocking antibodies were administered i.p. with an initial dose of 1mg/mouse 1 day before the tolerization (day 0); each mouse then received 0.5mg at day 3 and 6. IFN-α was blocked using 40μg/mouse per injection every 3 days during tolerization; IFN-λ2/3 blocking antibody (clone 244716, R&D Systems) or IgG isotype control (clone 141945, R&D Systems) were used. Tamoxifen (SIGMA) 10mg/ml was dissolved in warm corn oil in agitation for several hours at 37°C; mice were injected i.p. for 5 consecutive days at a concentration of 50mg/kg

### Hematoxylin-Eosin histology

Mice were euthanized 48 hours post intradermal OVA challenge, and ears collected in 10% PFA in PBS. Ears were embedded in paraffin, and 5 μm sections were stained with hematoxylin and eosin according to a standardized protocol.

### Antibody detection

Mice were bled at euthanization and serum samples were analyzed by ELISA in flat-bottom 96-well MaxiSorp microtiter plates (Nunc). Wells were coated with 5mg/ml OVA overnight in PBS at 4°C. Plates were washed and blocked with 0.1% bovine serum albumin (BSA) in PBS. Samples were then serially diluted in 0.1% BSA in PBS, and incubated overnight at 4°C. After washing, plates were incubated with 1:2000 anti mouse IgG1 Horseradish Peroxidase (HRP)-conjugated antibody, washed again and TMB was added as HRP substrate. Reaction was stopped with 0.16M H_2_SO_4_ and read at 450nm using a BioTek Synergy H1 microplate reader with Gen5 3.14 software. Antibody titers were determined as the reciprocal interpolated dilutions of the samples, giving rise to an absorbance on the linear part of the curve 0.2 above the background.

### Skin tissue processing

Single cell suspensions from ear skin were obtained as described previously (*16*). Briefly, mice were euthanized with CO_2_, and ear pinna skin was split into dorsal and ventral sheets and incubated in RPMI 1640 media supplemented with 2mM L-glutamine, 1mM sodium pyruvate, 1mM non-essential amino acids, 50μM β-mercaptoethanol, 20mM HEPES, 100U/ml of penicillin, 100mg/ml of streptomycin, 0.5mg/ml DNAse-I (Sigma-Aldrich) and 0.25mg/ml of Liberase TL (Roche) purified enzyme blend (Roche) for 1 hour and 45 min at 37°C/5% CO_2_. Digested ears were homogenized using the Medicon/Medimachine tissue homogenizer system (Becton Dickinson) and filtered through a 70μm cell strainer.

### Intestine and mesenteric lymph node tissue processing

Small intestine (SI), large intestine lamina propria and mesenteric lymph nodes (mLN) were collected and placed in complete cold RPMI 1640 media (supplemented as above). Mesenteric LNs were smashed through 70μm cell strainers, and for myeloid populations, digested in complete RPMI media with 0.5mg/ml DNAse I and 0.05mg/ml LiberaseTL for 20 minutes at 37°C/5% CO2 and then smashed through 70μm cell strainers. For SI and colon, Peyer’s patches and the mesenteric adipose tissue were removed. Tissues were cut opened, washed several times and cut into 1-2 cm segments. Tissues were treated with complete 3% FBS media containing 5mM EDTA and 0.145mg/ml dithiothreitol (DTT) for 20 minutes at 37°C/5% CO_2_ with constant stirring. Tissues were then shaken vigorously for 1min, three times, in 20mM HEPES, 2mM EDTA in RPMI 1640 media. For some experiments, epithelial fraction was collected at this step. After filtering, tissues were further digested with 10ml of media containing 500mg/ml DNAse I and 100mg/ml Liberase TL with continuous stirring at 37°C/5% CO_2_. Digested tissues were passed though 70-μm cell strainers, and leukocytes enriched by resuspension in 4ml of 37.5% Percoll and centrifugation at 400g for 5 minutes.

### Flow cytometry analysis

Single cell suspensions were incubated for 30 minutes at 4°C with surface marker fluorophore-conjugated antibodies in 2mM EDTA, 0.5% FBS in PBS, in presence of purified anti-mouse CD16/32 (FcBlock, clone 93). Dead cells were excluded using a LIVE/DEAD Fixable Blue Dead Cell Stain Kit (Invitrogen). Cells were fixed and permeabilized utilizing the Foxp3/Transcription Factor Staining Buffer Set (eBioscience) and stained with fluorophore-conjugated intracellular antibodies for at least 45 minutes at room temperature.

For TGF-β detection, single cell suspensions were culture *ex vivo* at 37°C for 2.5 hours in 10% FBS supplemented (as above) RPMI 1640 media, with 5 μg/mL Ionomycin (Sigma-Aldrich), 1:1000 dilution of GolgiPlug (BD Biosciences), and 50 ng/mL phorbol myristate acetate (PMA) (Sigma-Aldrich). Cells were intracellularly stained, as described above, with anti-LAP (TGF-β1) APC-conjugated antibody (Biolegend). Retinoic acid activity was analyzed by measuring RALDH activity in cells using the ALDEFLUOR kit (StemCell), following manufacturer’s instructions. LINE ORF1p was stained using an intracellular already conjugated specific antibody (Abcam), after fixation and permeabilization, for 1 hour at room temperature. For detection of MLV SU envelope protein, cells were incubated with hybridoma 83A25 supernatant to detect ecotropic and non-ecotropic ERV envelope protein (MLV Xmv45). Single cell suspensions were incubated for 1 hour at RT with the supernatant and FcBlock, followed by a biotinylated anti-rat IgG2A antibody (clone RG7/1.30, Biosciences) and streptavidin-BV650 (Biolegend).

### Gliadin diet and tetramer staining

Gliadin-containing diet (TD.00588) was purchased form Envigo Teklad Diets. Mice were fed for 7 days, and secondary lymphoid organs (SLOs: mLNs, Peyer’s patches, spleen and hepatic LNs) were obtained to analyze gliadin-specific cells as previously reported (*25*). As a control of the experiment, mice were fed control diet (TD.180916) not containing gliadin. Tetramers were obtained from the NIH Tetramer Core Facility: I-A(b) Wheat GLP 290-301 NVYIPPYCTIAP PE-labeled and I-A(b) Wheat GLP 290-301 NVYIPPYCTIAP APC-labeled. Secondary lymphoid organs (mLNs, Peyer’s patches, spleen and hepatic LNs) were pooled, smashed through 70μm cell strainers, and stained with tetramers for 1 hour at room temperature. Tetramer^+^ cells were enriched using anti-PE or anti-APC coupled magnetic beads (StemCell Technologies) and bead-bound cells were selected on magnetized columns (StemCell Technologies). Samples were then stained with surface and intracellular markers as described above. Normalized counts were calculated by experiment, dividing each sample by the average of control tetramer^+^ numbers in that experiment.

### 16sRNA library preparation, sequencing and analysis

DNA was extracted from fecal samples by using lysis matrix E tubes on the Precelleys tissue homogenizer and purified using a MagAttract PowerMicrobiome DNA/RNA EP Kit (Qiagen) on an automated liquid handling system (Eppendorf). Dual-indexing amplification and sequencing approach was taken to assess the composition of microbial communities from the given samples targeting the V4 hypervariable region of the 16S ribosomal RNA gene (*16S rRNA*), using primers 515F and 806R (*50*) and 105 ng of DNA as input. Libraries were then quantified using the KAPA library quantification kit (Roche), pooled at equimolar concentrations and sequenced on a MiSeq system (Illumina). 16S data was analyzed using Nephele’s DADA2 package (*51*, *52*). Taxonomy was assigned to each variant using the SILVA v138.1 database (*53*). Rooted phylogenetic tree was provided by Nephele’s standard output. Beta diversity metrics, including Weighted UniFrac and Unweighted UniFrac, were calculated using phyloseq package.

### T cells single cell RNA-seq

Small intestine CD4^+^ T cells with an enriched proportion of T_regs_ were sorted in a Sony MA900 sorter. Live (DAPI^-^) CD45^+^ CD11b^-^ CD11c^-^ CD19^-^ TCRψ8^-^ MHC-II^-^ CD64^-^ Ter119^-^ EpCAM^-^ TCRβ^+^ CD8^-^ CD4^+^ cells and Live (DAPI^-^) CD45^+^ CD11b^-^ CD11c^-^ CD19^-^ TCRψ8^-^ MHC-II^-^ CD64^-^ Ter119^-^ EpCAM^-^ TCRβ^+^ CD8^-^ CD4^+^ Foxp3-EGFP^+^ cells labeled with TotalSeqC hashtag antibodies (BioLegend) were sorted from 8-9 weeks old Foxp3-EGFP reporter mice treated or not with anti-reverse transcriptase treatment for 7 days (n=5 each condition). Sorted cells were pooled and combined in a 1:1 proportion (CD4^+^ T cells and CD4^+^ Foxp3^+^ Treg cells) and 40000 cells per lane (total of 3 lanes) were loaded to a Chromium Single Cell Controller (10X Genomics) to encapsulate cells into droplets. The gene expression (GEX), hashtag oligonucleotide (HTO) and VDJ (TCR) sequencing libraries were prepared using Chromium Single Cell 5’ v2 reagent kits following the manufacturer’s instructions and sequenced on an Illumina Nextseq2000 platform (NextSeq 1000/2000 P2 reagents). The initial processing of the expression data involved generating fastq files and count matrices using Cell Ranger v7.1.0 (*54*) (10X Genomics, Pleasanton, CA) that was run with the “Include introns=True” option and the mm10-2020-A transcriptome reference. The TCR data was processed with Cell Ranger v7.1.0 using a VDJ reference based on the IGMT database: The IGMT reference was generated with Cell Ranger v7.1.0 using the ‘fetch-imgt’ and ‘cellranger mkvdjref’ functions. Processing of the expression data was performed with the ‘cellranger multi’ and ‘cellranger aggr’ functions. After processing and aggregation, the estimated number of cells and mean reads per cell for each library were as follows: 54871 estimated number of cells, 23005 mean reads per cell (from 3 GEX libraries), 685 Median hashtag UMIs per Cell (across 3 HTO libraries), 38882 cells with a productive TRA and/or TRB V-J spanning sequence, 29108 of which contain both a productive V-J spanning TRA and TRB pair (from 3 VDJ libraries). The overall sequencing quality was medium to high in the 3 mRNA libraries: >93.5% of bases in the barcode and UMI regions had Q30 (99.99% inferred base call accuracy) quality score or above, whereas >81.7% of bases in the RNA reads had Q30 or above (mean 85.2%). Furthermore, across the 3 GEX libraries the median gene count per cell range was 1574-1876, the sequencing saturation ranged between 70.3-80.3%, and 72.1-76.9% of the reads mapped confidently to the transcriptome. Across the 3 TCR libraries, the mean used reads per cell was 4975-8956 (average 7593). Across the 3 hashtag (HTO) libraries the mean antibody reads usable per cell was 1044-2088 (average 1663).

The downstream analysis of the expression data was performed in 4.4.2, the Cell Ranger aggregation output (filtered_feature_bc_matrix directory) was loaded into Seurat version 5.1.0 (*55*) using the CreateSeuratObject function with min.cells=3. For filtering out low-quality cells, we established thresholds for gene and UMI counts (nCount_RNA < 30000, 200 < nFeature_RNA < 6000), and a maximum threshold for percentage mitochondrial content (percent.mt < 4) based on visual inspection of the corresponding metric distributions across all cells. Log-normalization (LogNormalize) of the RNA count data and centered log-ratio transformation (CLR) of the HTO count data were carried out using the NormalizeData function. For HTO demultiplexing cells into the individual mice and removing any doublets, HTODemux was run with a positive quantile = 0.99 and only cells identified as Singlets were kept. The top 2000 variable features were determined with the VariableFeatures function, expression data was scaled with ScaleData, and dimensionality reduction was performed with the RunPCA function on the variable features, followed by determining the dimensionality of the dataset on an elbow plot (ElbowPlot function), which was determined to be 35. Subsequently, FindNeighbors, FindClusters, and RunUMAP were run on 35 dimensions and a range of resolutions to determine the optimal resolution that captures biological diversity in the dataset, which was determined to be 1.25, resulting in 22 clusters. The clusters identified by the FindClusters function of Seurat (default Louvain clustering setting and 0.25 resolution) were annotated using cell type specific markers, and cluster specific differentially expressed genes. A small contaminating cluster comprised of Macrophages was removed.

For analysis of T regulatory cells, clusters annotated as Tregs were subset and FindVariableFeatures (nFeatures=2000) was run, data was scaled with ScaleData, RunPCA was run (npcs=100) and the top 30 dimensions were used to run FindNeighbors, FindClusters (with a range of resolutions and a final chosen resolution of 0.25), and RunUMAP. The resulting clusters were annotated using cell type specific markers, and cluster specific differentially expressed genes.

For analysis of T conventional (Foxp3^-^) cells, clusters not annotated as Tregs were subset and FindVariableFeatures (nFeatures=2000) was run, data was scaled with ScaleData, RunPCA was run (npcs=100) and the top 28 dimensions were used to run FindNeighbors, FindClusters (with a range of resolutions and a final chosen resolution of 0.6), and RunUMAP. The resulting clusters were annotated using cell type specific markers, and cluster specific differentially expressed genes.

For differentially expressed genes analysis per condition we used the Wilcoxon rank sum test implemented in the seurat function ‘FindMarkers’ with default parameters. The normalized counts (CPMs) were extracted from the seurat object and used for visualization along be corresponding adjusted p-values and log2 fold changes. Pathway enrichment analysis of upregulated or downregulated genes was performed using Metascape (*56*).

TCR repertoire sequencing data were analyzed with the scRepertoire package v2.2.1 (*57*) in R 4.4.2. Starting with the filtered contig annotations output table from ‘cellranger aggr’, the createHTOContigList, combineTCR, and combineExpression functions were used for combining the TCR data from each sample and for integration of the combined TCR data with the corresponding scRNAseq Seurat object. We utilized the amino acid sequence of the paired CDR3 regions of the paired alpha and beta chains as the clonotype definition. T cell1 without both an alpha and beta chain sequences were filtered for this analysis. T cell repertoire diversity was estimated using the D50 metric, which was calculated in R as the fraction of clonotypes, ordered by abundance, which account for 50 percent of total TCR sequences. Clonotype size was defined at the level of each biological replicate (Of all sequences: Rare: between 0% and 0.01%, Small: between 0.01% and 0.1%, Medium: between 0.1% and 1%, Large: between 1% and 10%, Hyperexpanded: between 10% and 100%). Cells with a putative iNKT TCR were defined as containing a TRAV11-TRAJ18 alpha chain V gene usage. TCR overlap between mice was defined with the same clonotype definition: at level of the CDR3 amino acid region for the combined alpha and beta chains, or a single alpha or beta chain if the other was not detected; computed in R and visualized using the package circlize (*58*).

### Dendritic cells single cell RNA-seq

Small intestine was divided in duodenum, jejunum and ileum and dendritic cells were sorted in a Sony MA900 sorter. Conventional dendritic cells (Live (DAPI^-^) CD45^+^ B220^-^ TCRβ^-^ TCRψ8^-^ NK1.1^-^ CD64^-^ CD11c^high^ MHC-II^high^) labeled with TotalSeqC hashtag antibodies (BioLegend) were sorted from 8-9 weeks old C57BL/6J mice treated or not with anti-reverse transcriptase treatment for 7 days (n=3 each condition). Sorted cells were pooled during sorting and 40000 cells per lane (total of 4 lanes) were loaded on a Chromium Single Cell Controller (10X Genomics) to encapsulate the cells into droplets. The gene expression (GEX) and hashtag oligonucleotide (HTO) sequencing libraries were prepared using Chromium Single Cell 5’ v2 reagent kits following the manufacturer’s instructions and sequenced on an Illumina Nextseq1000 platform (NextSeq 1000/2000 P2 reagents). The initial processing of the expression data involved generating fastq files and count matrices using Cell Ranger v7.1.0 (*54*) (10X Genomics, Pleasanton, CA) that was run with the “Include introns=True” option and the mm10-2020-A transcriptome reference. Processing of the expression data was performed with the ‘cellranger multi’ and ‘cellranger aggr’ functions. After processing and aggregation, the estimated number of cells and mean reads per cell for each library were as follows: 53233 estimated number of cells, 38590 mean reads per cell (from 4 GEX libraries), and 1839 Median hashtag UMIs per Cell (across 4 HTO libraries). The overall sequencing quality was high in the 4 mRNA libraries: >93.1% of bases in the barcode and UMI regions had Q30 (99.99% inferred base call accuracy) quality score or above, whereas >82.2% of bases in the RNA reads had Q30 or above. Furthermore, across the 4 GEX libraries the median gene count per cell range was 1879-1909 the sequencing saturation ranged between 69-76%, and 69-72% of the reads mapped confidently to the transcriptome. Across the 4 hashtag (HTO) libraries the mean antibody reads usable per cell was 15452-21358 (average 17218).

The downstream analysis of the expression data was performed in R 4.4.2, the Cell Ranger aggregation output (filtered_feature_bc_matrix directory) was loaded into Seurat version 4.4.0 (*55*) using the CreateSeuratObject function with min.cells=3. For filtering out low-quality cells, we established thresholds for gene and UMI counts (nCount_RNA < 40000, 250 < nFeature_RNA < 6000), and a maximum threshold for percentage mitochondrial content (percent.mt < 7.5) based on visual inspection of the corresponding metric distributions across all cells.

For HTO demultiplexing cells into individual mice and removing any doublets, we utilized cellhashR (*59*), using the following methods with a consensus threshold of 0.59: ‘bff_cluster’, ‘htodemux’, ‘bff_raw’, ‘multiseq’, ‘dropletutils’. The cell sample assignments were incorporate onto the seurat object and only cells identified as Singlets were kept.

Log-normalization (LogNormalize) of the RNA count data and centered log-ratio transformation (CLR) of the HTO count data were carried out using the NormalizeData function. The top 2000 variable features were determined with the VariableFeatures function, expression data was scaled with ScaleData, and dimensionality reduction was performed with the RunPCA function on the variable features, followed by determining the dimensionality of the dataset on an elbow plot (ElbowPlot function), which was determined to be 75. Subsequently, FindNeighbors, FindClusters, and RunUMAP were run on 75 dimensions and a range of resolutions to determine the optimal resolution that captures biological diversity in the dataset, which was determined to be 1.5. The clusters identified by the FindClusters function of Seurat (default Louvain clustering setting and 1.5 resolution) were annotated using cell type specific markers, and cluster specific differentially expressed genes. Contaminating clusters comprised of cell types other than dendritic cells were removed. On the remaining cells FindVariableFeatures (nFeatures=2000) was run, data was scaled with ScaleData, RunPCA was run (npcs=100) and the top 70 dimensions were used to run FindNeighbors, FindClusters (with a range of resolutions and a final chosen resolution of 0.5), and RunUMAP. Clusters were annotated using cell type specific markers, and cluster specific differentially expressed genes. A total of 13000, 4135 and 1045 cells were analyzed in duodenum, jejunum and ileum respectively.

Cells originating from the small intestinal samples were subset and FindVariableFeatures (nFeatures=2000) was run, data was scaled with ScaleData, RunPCA was run (npcs=100) and the top 60 dimensions were used to run FindNeighbors, FindClusters (with a range of resolutions and a final chosen resolution of 0.5), and RunUMAP. Clusters Were annotated using cell type specific markers, and cluster specific differentially expressed genes.

For differentially expressed genes analysis per small intestinal section per condition we used the Wilcoxon rank sum test implemented in the seurat function ‘FindMarkers’ with default parameters. The normalized counts (CPMs) were extracted from the seurat object and used for visualization along be corresponding adjusted p-values and log2 fold changes. Pathway enrichment analysis of upregulated or downregulated genes was performed using Metascape (*56*).

Ligand-receptor interactions were computed and visualized using the R package CellChat (v2.1.2) (*60*).

### Bulk RNA-seq

Epithelial cells and dendritic cells were sorted in a Sony MA900 sorter from duodenum, colon or mesenteric lymph nodes. Samples were staining with antibodies against CD45, B220, TCRβ, TCRψ8, NK1.1, CD64, CD11c, MHC-II, CD103, CD11b, EpCAM. Conventional dendritic cells type 1 (cDC1s: Live (DAPI^-^) CD45^+^ B220^-^ TCRβ^-^ TCRψ8^-^ NK1.1^-^ CD64^-^ CD11c^high^ MHC-II^high^ CD103^+^ CD11b^-^), CD103^+^ conventional dendritic cells type 2 (Live (DAPI^-^) CD103^+^ cDC2s: CD45^+^ B220^-^ TCRβ^-^ TCRψ8^-^ NK1.1^-^ CD64^-^ CD11c^high^ MHC-II^high^ CD103^+^ CD11b^-^) and epithelial cells (Live (DAPI^-^) CD45^-^ B220^-^ TCRβ^-^ TCRψ8^-^ NK1.1^-^ EpCAM^+^) were sorted directly into lysis buffer. RNA was purified using RNeasy Micro Kit (Qiagen) and libraries were prepared using Revelo High Sensitivity RNA-seq library preparation kit from Tecan according to manufacturer’s protocol. Bulk RNA-seq libraries were sequenced on a NextSeq1000. Endogenous retroelement expression was determined as previously described (*16*). Briefly, sequencing reads were mapped to the C57BL/6 mouse genome (GRCm38: mm10) and differential retroelement expression was calculated utilizing DESeq2 with homer’s getDifferentialExpression with default parameters (homer version 4.11). EREs with False Discover Rate (FDR) of < 0.05 and fold change > 2 were considered differentially expressed. For ERV analysis, homer’s analyzeRepeats with a custom ERV file (annotation provided by gEVE version 1.1 (*72*)) was utilized (for chromosomal location analysis), as well as with the parameter repeats (for locus analysis).

### cGAMP detection in gut explants

Duodenal gut explants were obtained as described elsewhere for colon (*36*). Briefly, intestinal content was flushed with ice-cold PBS several times, opened longitudinally and washed with ice-cold PBS. Pieces of approximately 0.5cm were weighted to normalize per mg tissue and placed in 48-well plate treated for tissue culture. Luminal side was placed facing up in DMEM/F12, 10%FBS, 1X penicillin/streptomycin media at kept at 37°C/5% CO_2_. Explants were treated or not with antiretroviral cocktail (Emtricitabine and Tenofovir) at 10μM. Supernatants were collected 24h later, centrifuged at 450g for 10min at 4°C and cGAMP was detected by ELISA (Cayman Chemical) according to manufacturer’s protocol.

### Statistical analysis

Statistical methods utilized are described in figure legends. Groups were compared using Prism software (version 10).

## Acknowledgements

We thank S. Mistry, Burns A, K. Beacht, E. Lewis, M. Abid, the NIAID Microbiome program sequencing platform, the NIAID animal facility and the NIAID Microbiome program Gnotobiotic animal facility for technical support; K. McCauley for data analysis support; the NIH Tetramer Core Facility (NIH Contract 75N93020D00005 and RRID:SCR_026557) for providing I-A(b) Wheat GLP 290-301 and control Tetramer; and all the Belkaid Laboratory members for helpful discussions and for providing constructive feedback on this project, particularly Dr. T. Farley for critical reading of the manuscript. Schematic figures were created with BioRender.

## Funding

This work was supported by the NIAID Division of Intramural Research. CAR, EA, VML and AT are supported by the Division of Intramural Research of NIAID (NIAID; 1ZIA-AI001115 and 1ZIA-AI001132); AT is in part supported by an EMBO Postdoctoral Fellowship (ALTF 1014-2021); CAR is the Lorraine W. Egan Fellow of the Damon Runyon Cancer Research Foundation (grant DRG-2496-23).

## Author contributions

Conceptualization: CAR, YB; Investigation: CAR, YB, EA, VML, SRK; Methodology: CAR, YB, EA, VML, SRK, AT, DY; Supervision: CAR, YB; Visualization: CAR, YB, EA, VML, SRK, CO; Writing – original draft: CAR, YB; Writing – review and editing: CAR, YB, EA, VML, SRK, AT, CO, DY.

## Competing interests

The authors declare no competing interests.

## Data and materials availability

RNAseq and scRNA-seq data were deposited into the Gene Expression Omnibus (GEO) data repository (GSE298407 for scRNA-seq of CD4^+^ T cell and T_regs_ in control and anti-RT conditions; GSE298496 for scRNA-seq of cDCs along the gut in control and anti-RT conditions; GSE297850 for bulk RNA-seq of epithelial cells and cDCs from SI, mLN and colon in control and anti-RT conditions; GSE297851 for bulk RNA-seq of epithelial cells and cDCs under different diets; GSE297852 for bulk RNA-seq of epithelial cells and cDCs in TLR-deficient mice). All data are available in the main text or the supplementary materials.

---

**Fig. S1.**
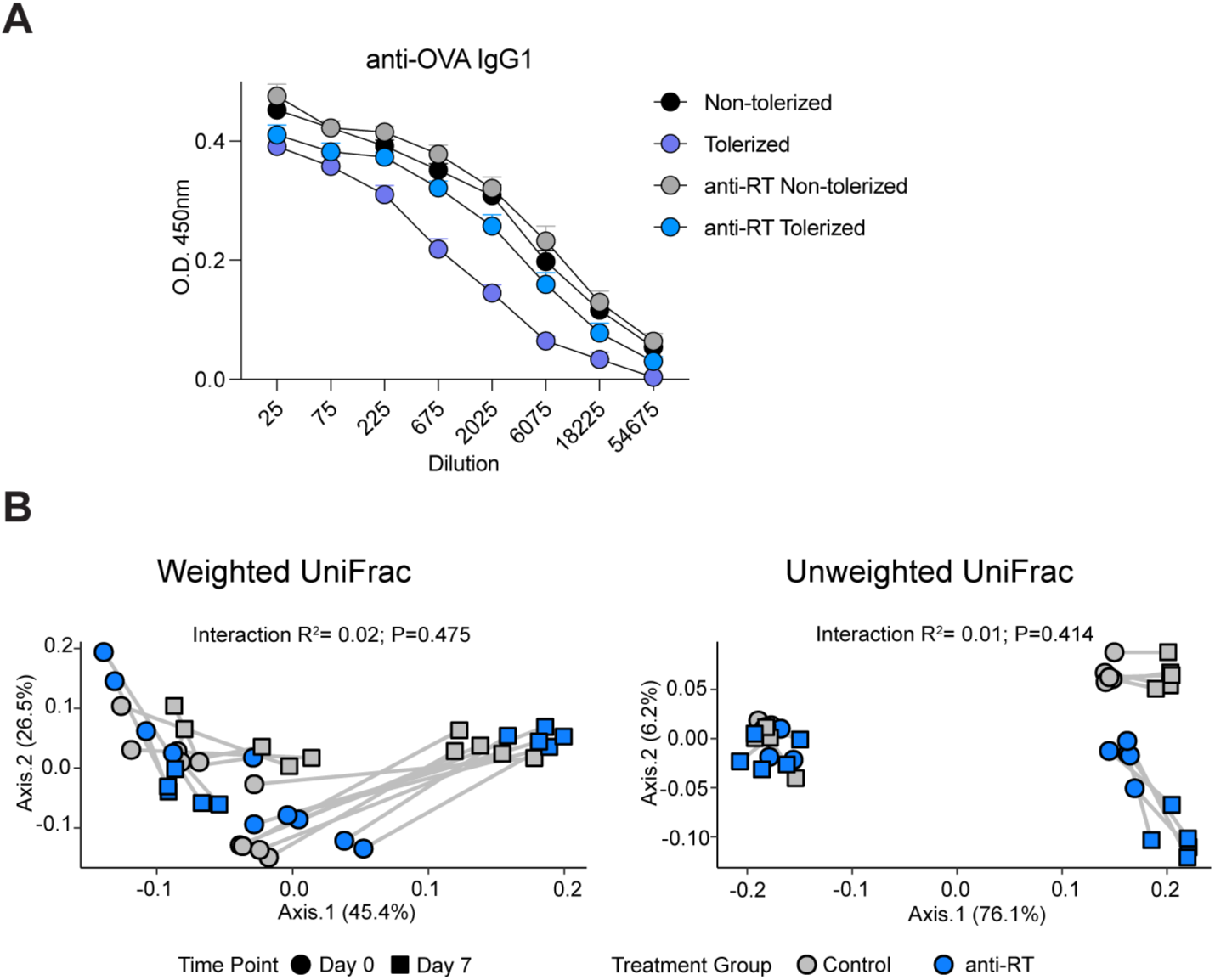
Retroelement reverse transcription effect on antibody production and microbiota composition. (**A**) OVA-specific IgG1 antibodies analyzed by ELISA in sera samples obtained 48h post ear challenge in the DTH model. Graph depicts 1 independent experiment representative of 3 independent experiments. (**B**) Paired analysis of unweighted and weighted UniFrac microbiota 16S profiles in control and anti-RT treated mice, before and after treatment. Percentages represent the variance explained by each PC.

**Fig. S2.**
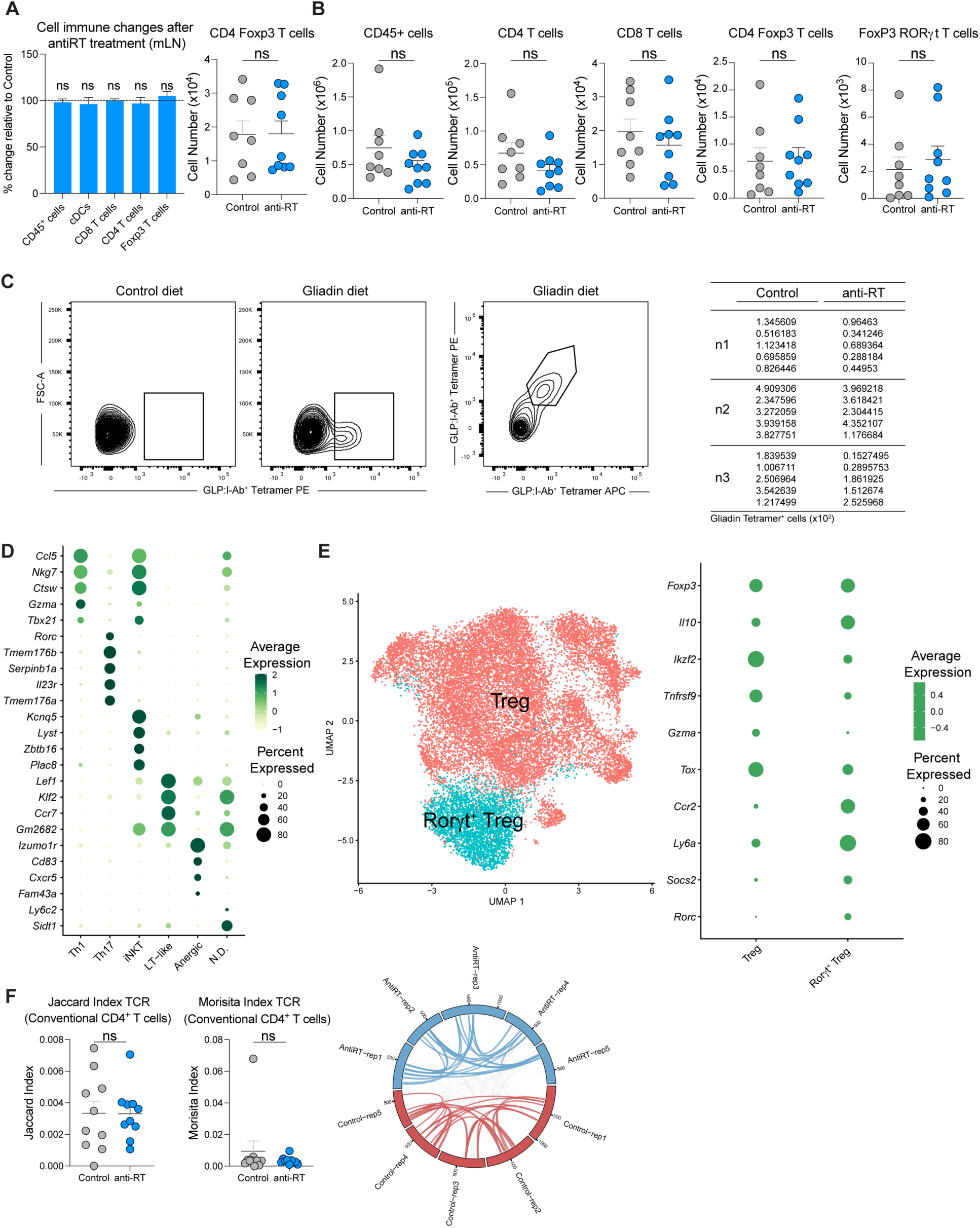
Inhibition of retroelement reverse transcription does not influence immune populations in mLN or colon, or TCR clonality in conventional CD4^+^ T cells. (**A**) (Left) Percentage change relative to control in absolute numbers of the indicated immune populations in mesenteric lymph nodes after 7 days of antiretroviral treatment (left). Live CD45^+^ CD90.2^+^ TCRβ^+^ CD4^+^ Foxp3^+^ cells absolute number quantification by flow cytometry analysis in mesenteric lymph nodes (right). (**B**) Live CD45^+^, Live CD45^+^ CD90.2^+^ TCRβ^+^ CD4^+^, Live CD45^+^ CD90.2^+^ TCRβ^+^ CD8^+^, Live CD45^+^ CD90.2^+^ TCRβ^+^ CD4^+^ Foxp3^+^ and Live CD45^+^ CD90.2^+^ TCRβ^+^ CD4^+^ Foxp3^+^ RorγT^+^ cells absolute number quantification by flow cytometry analysis in colon. (**C**) Representative contour plot of Glp:I-Ab tetramer^+^ CD4^+^ CD44^+^ Foxp3^+^ cells in secondary lymphoid organs (SLOs: mLNs, Peyer’s patches, spleen and hepatic LNs) and quantification analyzed by flow cytometry. (**D**) Dot plot of highly expressed genes in each cluster from CD4^+^ T cell single-cell RNAseq analysis, used for cluster annotation. (**E**) UMAP representation of Foxp3^+^ CD4^+^ cell clusters from single-cell RNAseq analysis (left). Dot plot of highly expressed genes in each cluster from Foxp3^+^ CD4^+^ T cells single-cell RNAseq analysis, used for cluster annotation. (**F**) Jaccard and Morisita indexes of conventional CD4^+^ T cell TCRs where each dot represents TCR overlap between mice under the same treatment (Left). Circos plot of conventional CD4^+^ T cell Receptor (TCR) analysis comparing control vs anti-reverse transcriptase treated mice. Each segment represents a mouse. Links between segments represent shared TCR between mice and colored links represent shared TCR between mice under the same treatment (Right). For flow cytometry analyses, data are representative of at least two independent experiments. Each dot represents an individual mouse. Numbers in flow plots indicate mean ± SEM. For (A), (B) and (F) two-tailed unpaired Student’s t-test or Mann-Whitney test were used; ns, not significant.

**Fig. S3.**
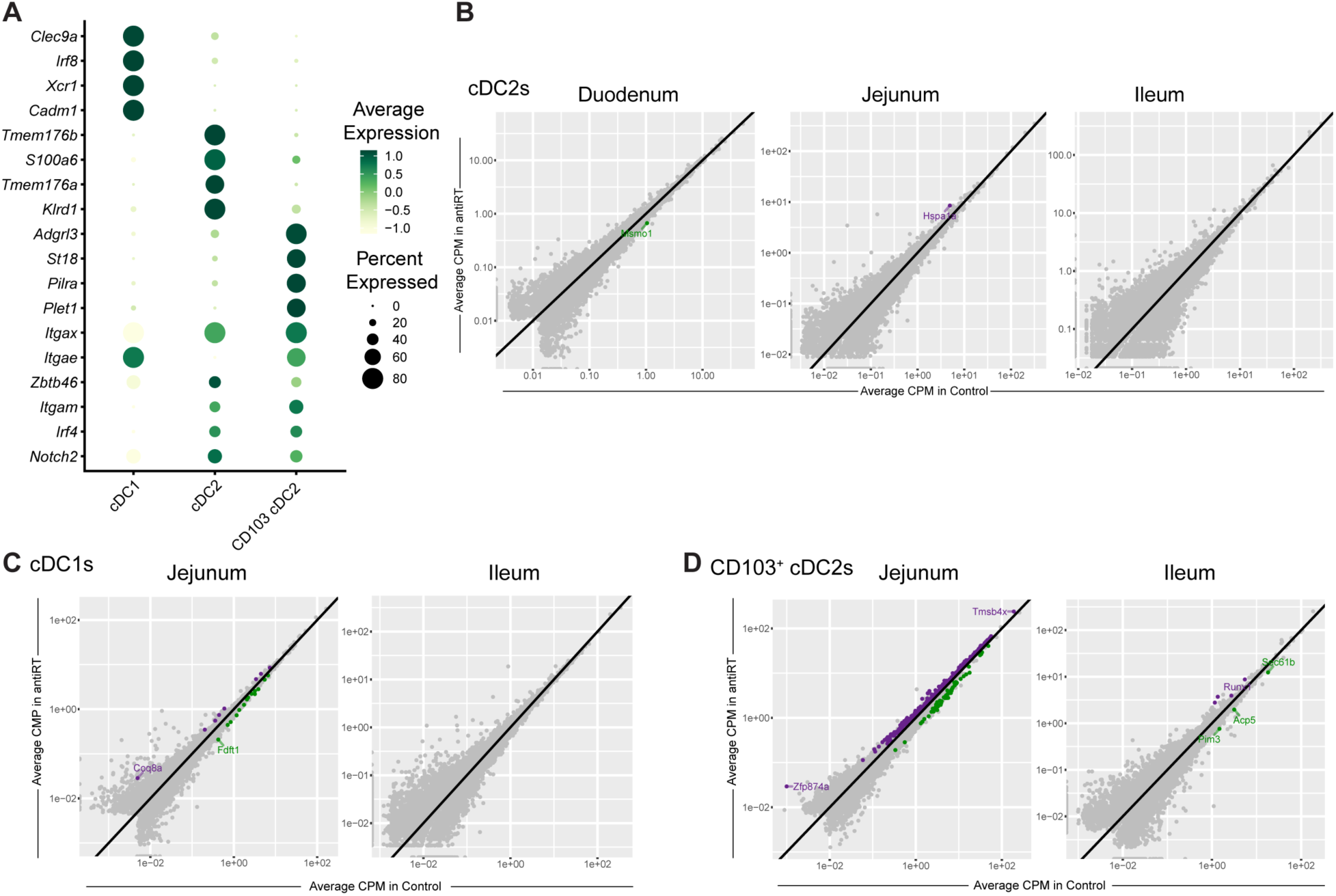
Retroelement activity does not modulate dendritic cell populations in the lower parts of the small intestine. (**A**) Dot plot of highly expressed genes in each cluster from MHC-II^high^ CDC11c^high^ cells single-cell RNAseq analysis, used for cluster annotation. (**B**) Scatter plots of DEGs in cDC2s from duodenum, jejunum and ileum. Purple denotes genes upregulated and green downregulated by anti-RT treatment. (**C**) Scatter plots of DEGs in cDC1s from jejunum and ileum after inhibition of reverse transcriptase treatment. Purple denotes genes upregulated and green downregulated by anti-RT treatment. (**D**) Scatter plots of DEGs in CD103^+^ cDC2s from jejunum and ileum after inhibition of reverse transcriptase treatment. Purple denotes genes upregulated and green downregulated by anti-RT treatment.

**Fig. S4.**
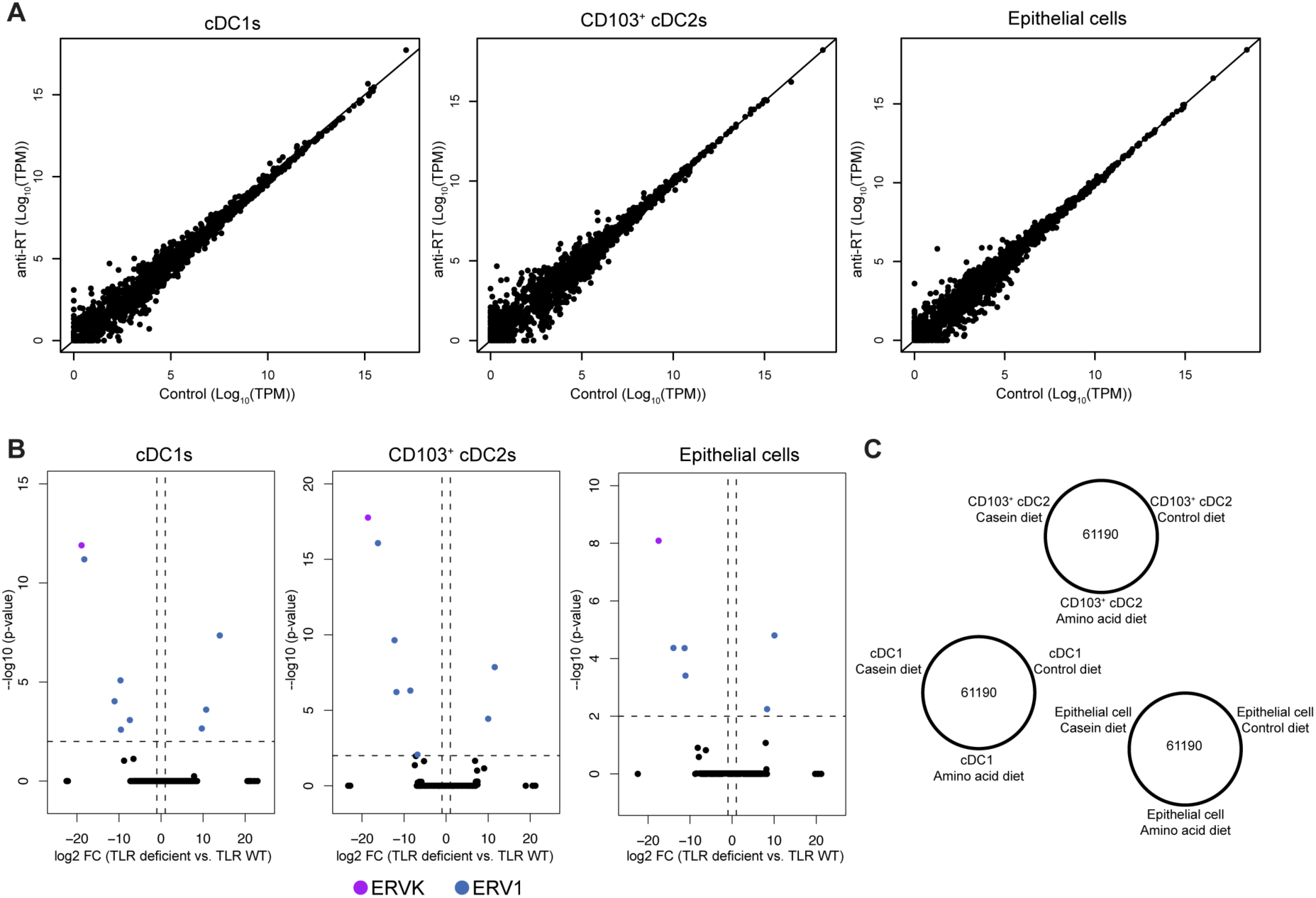
Endogenous retroelement expression in small intestine is independent of exogenous factors. (**A**) Retroelement expression analyzed by bulk RNA sequencing in cDC1s, CD103^+^ cDC2s and epithelial cells from duodenum, in control versus anti-RT treated mice. (**B**) Volcano plots of expressed retroelement loci from cDC1s (left), CD103^+^ cDC2s (middle) and epithelial cells (right) purified from the duodenum of WT or TLR deficient mice. (**C**) Venn diagram of expressed retroelements from bulk RNA-seq analysis. Mice were fed chow, casein, or amino acid (AA) diets since birth, and ERE expression was analyzed in gut epithelial cells, CD103^+^ cDC2s, and cDC1s from duodenum at 8 weeks old.

**Fig. S5.**
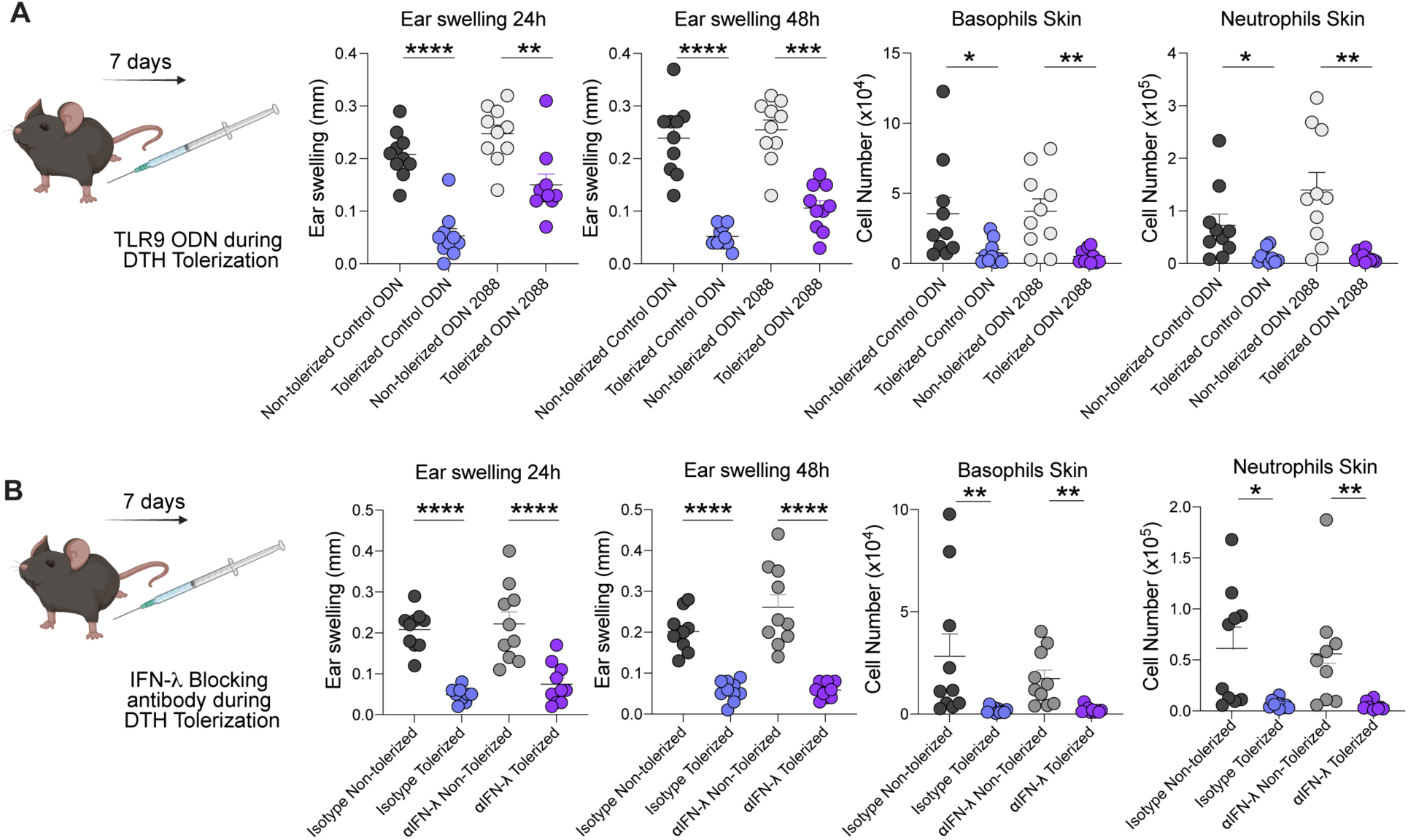
TLR9 and IFN-λ are not required for the establishment of oral tolerance. (**A**) TLR9 signaling pathway was temporally blocked during DTH tolerization using a specific inhibitory oligonucleotide (ODN 2088) or Control ODN. Ear swelling was analyzed after 24h and 48h, and infiltration of neutrophils and basophils were analyzed 48h later by flow cytometry in the skin. (**B**) Mice were treated with a blocking antibody for IFN lambda (2/3) or isotype control during DTH tolerization. Ear swelling was analyzed after 24h and 48h, and infiltration of neutrophils and basophils were analyzed 48h later by flow cytometry. Data are representative of at least two independent experiments. Each dot represents an individual mouse. For (A) and (B), one-way ANOVA with Tukey’s multiple comparisons test or Kruskal-Wallis with Dunn’s multiple comparisons test were used. Numbers in flow plots indicate mean ± SEM. * p < 0.05; ** p < 0.01; *** p < 0.001; **** p < 0.0001.

## Notes

### Competing Interest Statement

The authors have declared no competing interest.

## References

1. C. M. Warren, J. Jiang, R. S. Gupta, Epidemiology and Burden of Food Allergy. Curr Allergy Asthma Rep. 20 (2), 6 (2020).

2. D. Azzolino, L. Verdi, S. Perna, I. Baldassari, M. Cesari, T. Lucchi, Food allergies in older people: An emerging health problem. World Allergy Organization Journal. 17, 100967 (2024).

3. M. C. C. Canesso, T. B. R. Castro, S. Nakandakari-Higa, A. Lockhart, J. Luehr, J. Bortolatto, R. Parsa, D. Esterházy, M. Lyu, T.-T. Liu, K. M. Murphy, G. F. Sonnenberg, B. S. Reis, G. D. Victora, D. Mucida, Identification of antigen-presenting cell–T cell interactions driving immune responses to food. Science. 387 (6739):eado5088 (2025).

4. J. L. Coombes, K. R. R. Siddiqui, C. V. Arancibia-Cárcamo, J. Hall, C. M. Sun, Y. Belkaid, F. Powrie, A functionally specialized population of mucosal CD103+ DCs induces Foxp3+ regulatory T cells via a TGF-β- and retinoic acid-dependent mechanism. Journal of Experimental Medicine. 204 (2007).

5. C. M. Sun, J. A. Hall, R. B. Blank, N. Bouladoux, M. Oukka, J. R. Mora, Y. Belkaid, Small intestine lamina propria dendritic cells promote de novo generation of Foxp3 T reg cells via retinoic acid. Journal of Experimental Medicine. 204, 1775–1785 (2007).

6. G. Kassiotis, Endogenous Retroviruses and the Development of Cancer. The Journal of Immunology. 192 (2014).

7. J. D. Larouche, A. Trofimov, L. Hesnard, G. Ehx, Q. Zhao, K. Vincent, C. Durette, P. Gendron, J. P. Laverdure, É. Bonneil, C. Côté, S. Lemieux, P. Thibault, C. Perreault, Widespread and tissue-specific expression of endogenous retroelements in human somatic tissues. Genome Med. 12 (2020).

8. G. Kassiotis, The Immunological Conundrum of Endogenous Retroelements. Annu Rev Immunol. 15 (2024).

9. M. Friedli, D. Trono, The Developmental Control of Transposable Elements and the Evolution of Higher Species. Annu Rev Cell Dev Biol. 31, 429–451 (2015).

10. N. Grandi, E. Tramontano, Human endogenous retroviruses are ancient acquired elements still shaping innate immune responses. Front Immunol. 9, 2039 (2018).

11. S. Srinivasachar Badarinarayan, D. Sauter, Switching Sides: How Endogenous Retroviruses Protect Us from Viral Infections. J Virol. 95 (2021).

12. C. Lavialle, G. Cornelis, A. Dupressoir, C. Esnault, O. Heidmann, C. Vernochet, T. Heidmann, Paleovirology of “syncytins”, retroviral env genes exapted for a role in placentation. Phil Trans R Soc B. 368, 20120507 (2013).

13. M. Dolci, C. Favero, W. Toumi, E. Favi, L. Tarantini, L. Signorini, G. Basile, V. Bollati, S. D’Alessandro, P. Bagnoli, P. Ferrante, S. Delbue, Human Endogenous Retroviruses Long Terminal Repeat Methylation, Transcription, and Protein Expression in Human Colon Cancer. Front Oncol. 10 (2020).

14. P. Küry, A. Nath, A. Créange, A. Dolei, P. Marche, J. Gold, G. Giovannoni, H. P. Hartung, H. Perron, Human Endogenous Retroviruses in Neurological Diseases. Trends in Molecular Medicine. 24, 4 (2018).

15. M. C. Steiner, J. L. Marston, L. P. Iñiguez, M. L. Bendall, K. B. Chiappinelli, D. F. Nixon, K. A. Crandall, Locus-specific characterization of human endogenous retrovirus expression in prostate, breast, and colon cancers. Cancer Res. 81 (2021).

16. D. S. Lima-Junior, S. R. Krishnamurthy, N. Bouladoux, N. Collins, S. J. Han, E. Y. Chen, M. G. Constantinides, V. M. Link, A. I. Lim, M. Enamorado, C. Cataisson, L. Gil, I. Rao, T. K. Farley, G. Koroleva, J. Attig, S. H. Yuspa, M. A. Fischbach, G. Kassiotis, Y. Belkaid, Endogenous retroviruses promote homeostatic and inflammatory responses to the microbiota. Cell. 184 (2021).

17. F. Fueyo-González, M. McGinty, M. Ningoo, L. Anderson, C. Cantarelli, Andrea Angeletti, M. Demir, I. Llaudó, C. Purroy, N. Marjanovic, D. Heja, S. C. Sealfon, P. S. Heeger, P. Cravedi, M. Fribourg, Interferon-β acts directly on T cells to prolong allograft survival by enhancing regulatory T cell induction through Foxp3 acetylation. Immunity. 55 (2022).

18. A. V. Ayala, C. Y. Hsu, R. E. Oles, K. Matsuo, L. R. Loomis, E. Buzun, M. C. Terrazas, R. R. Gerner, H. H. Lu, S. Kim, Z. Zhang, J. H. Park, P. Rivaud, M. Thomson, L. F. Lu, B. Min, H. Chu, Commensal bacteria promote type I interferon signaling to maintain immune tolerance in mice. J Exp Med. 221 (2024).

19. X. Zhang, L. Izikson, L. Liu, H. L. Weiner. Activation of CD25 CD4 Regulatory T Cells by Oral Antigen Administration. J Immunol. 167 (8), 4245–4253 (2001).

20. D. Mucida, N. Kutchukhidze, A. Erazo, M. Russo, J. J. Lafaille, M. A. Curotto De Lafaille, Oral tolerance in the absence of naturally occurring Tregs. Journal of Clinical Investigation. 115 (2005).

21. B. H. Yang, S. Hagemann, P. Mamareli, U. Lauer, U. Hoffmann, M. Beckstette, L. Föhse, I. Prinz, J. Pezoldt, S. Suerbaum, T. Sparwasser, A. Hamann, S. Floess, J. Huehn, M. Lochner, Foxp3+ T cells expressing RORγt represent a stable regulatory T-cell effector lineage with enhanced suppressive capacity during intestinal inflammation. Mucosal Immunol. 9 (2016).

22. K. A. Knoop, K. G. McDonald, C. S. Hsieh, P. I. Tarr, R. D. Newberry, Regulatory T Cells Developing Peri-Weaning Are Continually Required to Restrain Th2 Systemic Responses Later in Life. Front Immunol. 11 (2021).

23. Y. Wang, M. A. Su, Y. Y. Wan, An Essential Role of the Transcription Factor GATA-3 for the Function of Regulatory T Cells. Immunity. 35, 337–348 (2011).

24. U. Hadis, B. Wahl, O. Schulz, M. Hardtke-Wolenski, A. Schippers, N. Wagner, W. Müller, T. Sparwasser, R. Förster, O. Pabst, Intestinal Tolerance Requires Gut Homing and Expansion of FoxP3+ Regulatory T Cells in the Lamina Propria. Immunity. 34, 237–246 (2011).

25. S.-W. Hong, P. D. Krueger, K. C. Osum, T. Dileepan, A. Herman, D. L. Mueller, M. K. Jenkins, Immune tolerance of food is mediated by layers of CD4+ T cell dysfunction. Nature. 607 (7920), 762–768 (2022).

26. A. Lockhart, A. Reed, T. Rezende de Castro, C. Herman, M. C. Campos Canesso, D. Mucida, Dietary protein shapes the profile and repertoire of intestinal CD4+ T cells. J Exp Med. 220 (2023).

27. E. Ronin, M. L. Di Ricco, R. Vallion, J. Divoux, H. K. Kwon, S. Grégoire, D. Collares, A. Rouers, V. Baud, C. Benoist, B. L. Salomon, The nf-κb rela transcription factor is critical for regulatory t cell activation and stability. Front Immunol 10 (2019).

28. R. J. Miragaia, T. Gomes, A. Chomka, L. Jardine, A. Riedel, A. N. Hegazy, N. Whibley, A. Tucci, X. Chen, I. Lindeman, G. Emerton, T. Krausgruber, J. Shields, M. Haniffa, F. Powrie, S. A. Teichmann, Single-Cell Transcriptomics of Regulatory T Cells Reveals Trajectories of Tissue Adaptation. Immunity 50, 493–504.e7 (2019).

29. A. Vasanthakumar, Y. Liao, P. Teh, M. F. Pascutti, A. E. Oja, A. L. Garnham, R. Gloury, J. C. Tempany, T. Sidwell, E. Cuadrado, P. Tuijnenburg, T. W. Kuijpers, N. Lalaoui, L. A. Mielke, V. L. Bryant, P. D. Hodgkin, J. Silke, G. K. Smyth, M. A. Nolte, W. Shi, A. Kallies, The TNF Receptor Superfamily-NF-κB Axis Is Critical to Maintain Effector Regulatory T Cells in Lymphoid and Non-lymphoid Tissues. Cell Rep. 20, 2906–2920 (2017).

30. E. Jaensson-Gyllenbäck, K. Kotarsky, F. Zapata, E. K. Persson, T. E. Gundersen, R. Blomhoff, W. W. Agace, Bile retinoids imprint intestinal CD103+ dendritic cells with the ability to generate gut-tropic T cells. Mucosal Immunol. 4 (2011).

31. E. Jaensson, H. Uronen-Hansson, O. Pabst, B. Eksteen, J. Tian, J. L. Coombes, P. L. Berg, T. Davidsson, F. Powrie, B. Johansson-Lindbom, W. W. Agace, Small intestinal CD103+ dendritic cells display unique functional properties that are conserved between mice and humans. J Exp Med. 205 (2008).

32. G. R. Young, B. Mavrommatis, G. Kassiotis, Microarray analysis reveals global modulation of endogenous retroelement transcription by microbes. Retrovirology. 11, 59 (2014).

33. K. E. Sivick, N. Arpaia, G. L. Reiner, B. L. Lee, B. R. Russell, G. M. Barton, Toll-like receptor-deficient mice reveal how innate immune signaling influences salmonella virulence strategies. Cell Host Microbe. 15, 203–213 (2014).

34. P. Yu, W. Lübben, H. Slomka, J. Gebler, M. Konert, C. Cai, L. Neubrandt, O. Prazeres da Costa, S. Paul, S. Dehnert, K. Döhne, M. Thanisch, S. Storsberg, L. Wiegand, A. Kaufmann, M. Nain, L. Quintanilla-Martinez, S. Bettio, B. Schnierle, L. Kolesnikova, S. Becker, M. Schnare, S. Bauer, Nucleic Acid-Sensing Toll-like Receptors Are Essential for the Control of Endogenous Retrovirus Viremia and ERV-Induced Tumors. Immunity. 37, 867–879 (2012).

35. R. E. Rigby, L. M. Webb, K. J. Mackenzie, Y. Li, A. Leitch, M. A. M. Reijns, R. J. Lundie, A. Revuelta, D. J. Davidson, S. Diebold, Y. Modis, A. S. MacDonald, A. P. Jackson, RNA:DNA hybrids are a novel molecular pattern sensed by TLR9. EMBO Journal. 33, 542–558 (2014).

36. R. T. Mertens, A. Misra, P. Xiao, S. Baek, J. M. Rone, D. Mangani, K. N. Sivanathan, A. S. Arojojoye, S. G. Awuah, I. Lee, G. P. Shi, B. Petrova, J. R. Brook, A. C. Anderson, R. A. Flavell, N. Kanarek, M. Hemberg, R. Nowarski, A metabolic switch orchestrated by IL-18 and the cyclic dinucleotide cGAMP programs intestinal tolerance. Immunity. 57 (9), 2077–2094 (2024).

37. N. Jeremiah, H. Ferran, K. Antoniadou, K. De Azevedo, J. Nikolic, M. Maurin, P. Benaroch, N. Manel, RELA tunes innate-like interferon I/III responses in human T cells. J Exp Med. 220 (2023).

38. J. Gutierrez-Merino, B. Isla, T. Combes, F. Martinez-Estrada, C. Maluquer De Motes, Beneficial bacteria activate type-I interferon production via the intracellular cytosolic sensors STING and MAVS. Gut Microbes. 11 (2020).

39. G. Kassiotis, J. P. Stoye. Immune responses to endogenous retroelements: Taking the bad with the good. Nat Rev Immunol. 16 (4), 207–19 (2016).

40. P. E. Bonté, Y. A. Arribas, A. Merlotti, M. Carrascal, J. V. Zhang, E. Zueva, Z. A. Binder, C. Alanio, C. Goudot, S. Amigorena. Single-cell RNA-seq-based proteogenomics identifies glioblastoma-specific transposable elements encoding HLA-I-presented peptides. Cell Rep. 39 (2022).

41. S. F. Erttmann, P. Swacha, K. M. Aung, B. Brindefalk, H. Jiang, A. Härtlova, B. E. Uhlin, S. N. Wai, N. O. Gekara. The gut microbiota prime systemic antiviral immunity via the cGAS-STING-IFN-I axis. Immunity. 55, 847–861.e10 (2022).

42. Y. Yang, L. Wang, I. Peugnet-González, D. Parada-Venegas, G. Dijkstra, K. N. Faber. cGAS-STING signaling pathway in intestinal homeostasis and diseases. Front Immunol. 14, 14, 1239142 (2023).

43. C. A. Stewart, H. Metheny, N. Iida, L. Smith, M. Hanson, F. Steinhagen, R. M. Leighty, A. Roers, C. L. Karp, W. Müller, G. Trinchieri. Interferon-dependent IL-10 production by Tregs limits tumor Th17 inflammation. Journal of Clinical Investigation. 123, 4859–4874 (2013).

44. P. H. Brigleb, E. Kouame, K. L. Fiske, G. M. Taylor, K. Urbanek, L. Medina Sanchez, R. Hinterleitner, B. Jabri, T. S. Dermody. NK cells contribute to reovirus-induced IFN responses and loss of tolerance to dietary antigen. JCI Insight. 7 (16):e159823 (2022).

45. R. Bouziat, R. Hinterleitner, J. J. Brown, J. E. Stencel-Baerenwald, M. Ikizler, T. Mayassi, M. Meisel, S. M. Kim, V. Discepolo, A. J. Pruijssers, J. D. Ernest, J. A. Iskarpatyoti, L. M. M. Costes, I. Lawrence, B. A. Palanski, M. Varma, M. A. Zurenski, S. Khomandiak, N. Mcallister, P. Aravamudhan, K. W. Boehme, F. Hu, J. N. Samsom, H.-C. Reinecker, S. S. Kupfer, S. Guandalini, C. E. Semrad, V. Abadie, C. Khosla, L. B. Barreiro, R. J. Xavier. Reovirus infection triggers inflammatory responses to dietary antigens and development of celiac disease. Science. 356 (6333), 44–50 (2017).

46. D. M. Da Fonseca, T. W. Hand, S. J. Han, M. Y. Gerner, A. G. Zaretsky, A. L. Byrd, O. J. Harrison, A. M. Ortiz, M. Quinones, G. Trinchieri, J. M. Brenchley, I. E. Brodsky, R. N. Germain, G. J. Randolph, Y. Belkaid, Microbiota-Dependent Sequelae of Acute Infection Compromise Tissue-Specific Immunity. Cell. 163, 354–366 (2015).

47. G. Oldenhove, N. Bouladoux, E. A. Wohlfert, J. A. Hall, D. Chou, L. Dos santos, S. O’Brien, R. Blank, E. Lamb, S. Natarajan, R. Kastenmayer, C. Hunter, M. E. Grigg, Y. Belkaid, Decrease of Foxp3+ Treg Cell Number and Acquisition of Effector Cell Phenotype during Lethal Infection. Immunity. 31, 772–786 (2009).

48. K. D. Jackson, M. P. H. Lajeana, D. Howie, C. H. E. S. Lara, J. Akinbami. Trends in Allergic Conditions among children: United States, 1997-2011. NCHS Data Brief. 121 (2013).

49. B. Zablotsky, L. I. Black, L. J. Akinbami. Diagnosed Allergic Conditions in Children Aged 0-17 Years: United States, 2021. NHCS Data Brief. 459 (2023)

50. J. G. Caporaso, C. L. Lauber, W. A. Walters, D. Berg-Lyons, C. A. Lozupone, P. J. Turnbaugh, N. Fierer, R. Knight, Global patterns of 16S rRNA diversity at a depth of millions of sequences per sample. Proc Natl Acad Sci U S A. 108, 4516–4522 (2011).

51. B. J. Callahan, P. J. McMurdie, M. J. Rosen, A. W. Han, A. J. A. Johnson, S. P. Holmes, DADA2: High-resolution sample inference from Illumina amplicon data. Nat Methods. 13, 581–583 (2016).

52. N. Weber, D. Liou, J. Dommer, P. Macmenamin, M. Quiñones, I. Misner, A. J. Oler, J. Wan, L. Kim, M. Coakley McCarthy, S. Ezeji, K. Noble, D. E. Hurt, Nephele: A cloud platform for simplified, standardized and reproducible microbiome data analysis. Bioinformatics. 34, 1411–1413 (2018).

53. C. Quast, E. Pruesse, P. Yilmaz, J. Gerken, T. Schweer, P. Yarza, J. Peplies, F. O. Glöckner, The SILVA ribosomal RNA gene database project: Improved data processing and web-based tools. Nucleic Acids Res. 41 (2013).

54. G. X. Y. Zheng, J. M. Terry, P. Belgrader, P. Ryvkin, Z. W. Bent, R. Wilson, S. B. Ziraldo, T. D. Wheeler, G. P. McDermott, J. Zhu, M. T. Gregory, J. Shuga, L. Montesclaros, J. G. Underwood, D. A. Masquelier, S. Y. Nishimura, M. Schnall-Levin, P. W. Wyatt, C. M. Hindson, R. Bharadwaj, A. Wong, K. D. Ness, L. W. Beppu, H. J. Deeg, C. McFarland, K. R. Loeb, W. J. Valente, N. G. Ericson, E. A. Stevens, J. P. Radich, T. S. Mikkelsen, B. J. Hindson, J. H. Bielas, Massively parallel digital transcriptional profiling of single cells. Nat Commun. 8 (2017).

55. Y. Hao, S. Hao, E. Andersen-Nissen, W. M. Mauck, S. Zheng, A. Butler, M. J. Lee, A. J. Wilk, C. Darby, M. Zager, P. Hoffman, M. Stoeckius, E. Papalexi, E. P. Mimitou, J. Jain, A. Srivastava, T. Stuart, L. M. Fleming, B. Yeung, A. J. Rogers, J. M. McElrath, C. A. Blish, R. Gottardo, P. Smibert, R. Satija, Integrated analysis of multimodal single-cell data. Cell. 184, 3573–3587.e29 (2021).

56. Y. Zhou, B. Zhou, L. Pache, M. Chang, A. H. Khodabakhshi, O. Tanaseichuk, C. Benner, S. K. Chanda, Metascape provides a biologist-oriented resource for the analysis of systems-level datasets. Nat Commun. 10 (2019).

57. Q. Yang, K. R. Safina, N. Borcherding. scRepertoire 2: Enhanced and Efficient Toolkit for Single-Cell Immune Profiling. [Preprint] (2024). 10.1101/2024.12.31.630854.

58. Z. Gu, L. Gu, R. Eils, M. Schlesner, B. Brors, Circlize implements and enhances circular visualization in R. Bioinformatics. 30, 2811–2812 (2014).

59. G. J. Boggy, G. W. Mcelfresh, E. Mahyari, A. B. Ventura, S. G. Hansen, L. J. Picker, B. N. Bimber. BFF and cellhashR: analysis tools for accurate demultiplexing of cell hashing data. Bioinformatics. 38 (10), 2791–2801 (2022).

60. S. Jin, C. F. Guerrero-Juarez, L. Zhang, I. Chang, R. Ramos, C. H. Kuan, P. Myung, M. V. Plikus, Q. Nie, Inference and analysis of cell-cell communication using CellChat. Nat Commun. 12 (2021).

